# Spatial and Temporal Organization of Odor-evoked Responses in a Fly Olfactory Circuit: Inputs, Outputs and Idiosyncrasies

**DOI:** 10.1101/2020.10.21.349688

**Authors:** Haoyang Rong, Lijun Zhang, Cody Greer, Yehuda Ben-Shahar, Tim Holy, Barani Raman

## Abstract

How is sensory information routed through different types of neurons within a circuit, and do equivalent circuits in different individuals follow similar organizational principles? We examined this issue in the fruit fly olfactory system. Odor-evoked signals from sensory neurons (ORNs) triggered neural responses that were patterned over space and time in cholinergic ePNs and GABAergic iPNs within the antennal lobe. The dendritic-axonal (I/O) response mapping was complex and diverse, and axonal organization was region-specific (mushroom body vs. lateral horn). In the lateral horn, feed-forward excitatory and inhibitory axonal projections matched ‘odor tuning’ in a stereotyped, dorsal-lateral locus, but mismatched in most other locations. In the temporal dimension, ORN, ePN and iPN odor-evoked responses had similar encoding features, such as information refinement over time and divergent ON and OFF responses. Notably, analogous spatial and temporal coding principles were observed in all flies, and the latter emerged from idiosyncratic neural processing approaches.

**Highlights:**

- Consistency and idiosyncrasy both exist in ORN, ePN and iPN functional maps
- Signal transformations between different ePN compartments are complex and diverse
- Temporal decorrelation between stimuli happens in all three neuronal populations
- OFF responses that are orthogonal to ON responses emerge after odor termination

## Introduction

Most neuronal networks consist of many sub-types of neurons that interact through different microcircuits and actively reorganize the information they receive. To fully understand the information processing carried out, at a bare minimum three pieces of information are essential. First, it is necessary to understand the input received by the network. Second, to understand what computations arise from which microcircuit, it is necessary to follow this input signal as it propagates from one processing compartment to the next. And, third, it is necessary to understand how different neuronal sub-types that are present in these circuits contribute to the information processing. An additional layer of investigation could be added by comparing how information is represented by equivalent circuits in different individuals. This would allow us to understand what are the generic rules of signal processing and information transformation, and help identify any idiosyncratic features that may be utilized in different individuals. Understanding such idiosyncrasies in neural encoding can arguably help us better understand a source of variance in behavioral outcomes observed across individuals. Here, we dissect how odor signals are organized and processed as it propagates through the fruit fly (*Drosophila melanogaster*) antennal lobe neural network.

In the fruit fly olfactory system, vapors from volatile chemicals are transduced into neural responses by olfactory receptor neurons (ORN) present in the antenna that then transmit this information to a region called the antennal lobe (analogous to the mammalian olfactory bulb). The ORNs of the same type, i.e. expressing the same receptor–co-receptor gene combination, send their axons to either one or two spherical structure of neuropil called glomeruli in the antennal lobe (Couto et al., 2005; Fishilevich and Vosshall, 2005). The ORN activity drives responses in three major types of neurons in the antennal lobe: GABAergic local neurons (LNs), cholinergic projection neurons (excitatory PNs or ePNs) and GABAergic projection neurons (inhibitory PNs or iPNs). The local neurons are diverse (Chou et al., 2010), and play important roles in how sensory signals are processed within the antennal lobe (Olsen and Wilson, 2008; Yaksi and Wilson, 2010). However, LNs do not send their processes outside the antennal lobe, and thus only the activity ePNs and iPNs constitute the outputs from this olfactory neuronal network.

Notably, the ePNs and iPNs differ in how they receive inputs and transmit their output. The ePN dendrites innervate a single glomerulus and therefore receive input from a single ORN type (Couto et al., 2005). The ePNs project their axons onto both mushroom body (a center associated with learning and memory (Debelle and Heisenberg, 1994; Heisenberg et al., 1985)) and lateral horn (a region with putative role in driving innate behavior (Gupta and Stopfer, 2012; Heimbeck et al., 2001). In contrast, iPNs dendrites are multi-glomerular and therefore integrate information distributed across several different ORN types. The iPN axons are also exclusively sent to the lateral horns. The ePNs and iPNs can influence each other’s activity through chemical synapses (Shimizu and Stopfer, 2017). While the importance of the ePN and iPN activity for odor recognition is well established (Ahsan et al., 2017; Parnas et al., 2013; Strutz et al., 2014), how the ePN and iPN activities are spatially organized and patterned over time to facilitate odor recognition remains poorly understood.

In this study, we used an *in vivo*, light-sheet, volumetric, calcium-imaging technique to examine this issue with high spatial and temporal resolution. We monitored the odor-evoked signals at the ORN axons entering the antennal lobe (input), the responses they drive in ePNs dendrites located within the antennal lobe, and ePN and iPNs axons (output) entering mushroom body calyx and lateral horn (iPNs only project to the latter). Using this approach, we examined how odorants-evoked responses are patterned over space and time in each of these neural population. We examined the functional mapping between dendritic and axonal compartments to understand the antennal lobe input-output relationships, and how feed-forward excitation and feed-forward inhibition converge onto lateral horn. Lastly, comparison across flies helped understand generic odor coding principles and how they might arise from idiosyncratic processing mechanisms utilized within the antennal lobe network.

## Results

### Light-sheeting imaging of odor evoked neural activity

We used a custom-built light-sheet imaging setup (Greer and Holy, 2019) to monitor calcium signals (GCamp6f) from olfactory sensory neurons expressing the *orco* co-receptor (ORNs), and their two downstream targets excitatory GH146 projection neurons (ePNs) and inhibitory Mz699 projection neurons (iPNs) (**Figure 1A - C**). In each fly, one of these three neural population was labeled, and neural responses from all optical planes was near-simultaneously recorded (see **Methods**; **Figure 1D**). While the axonal outputs alone were monitored for ORNs and iPNs (as GCamp6f expression levels were weak in the antennal lobe for the Mz699 line), both dendritic and axonal calcium signals were monitored for ePNs (GH146 line). This approach allowed us to relate the dendritic inputs in the antennal lobe with the functional signals reaching the two downstream targets: mushroom body calyces and lateral horns.

**Figure 1:**
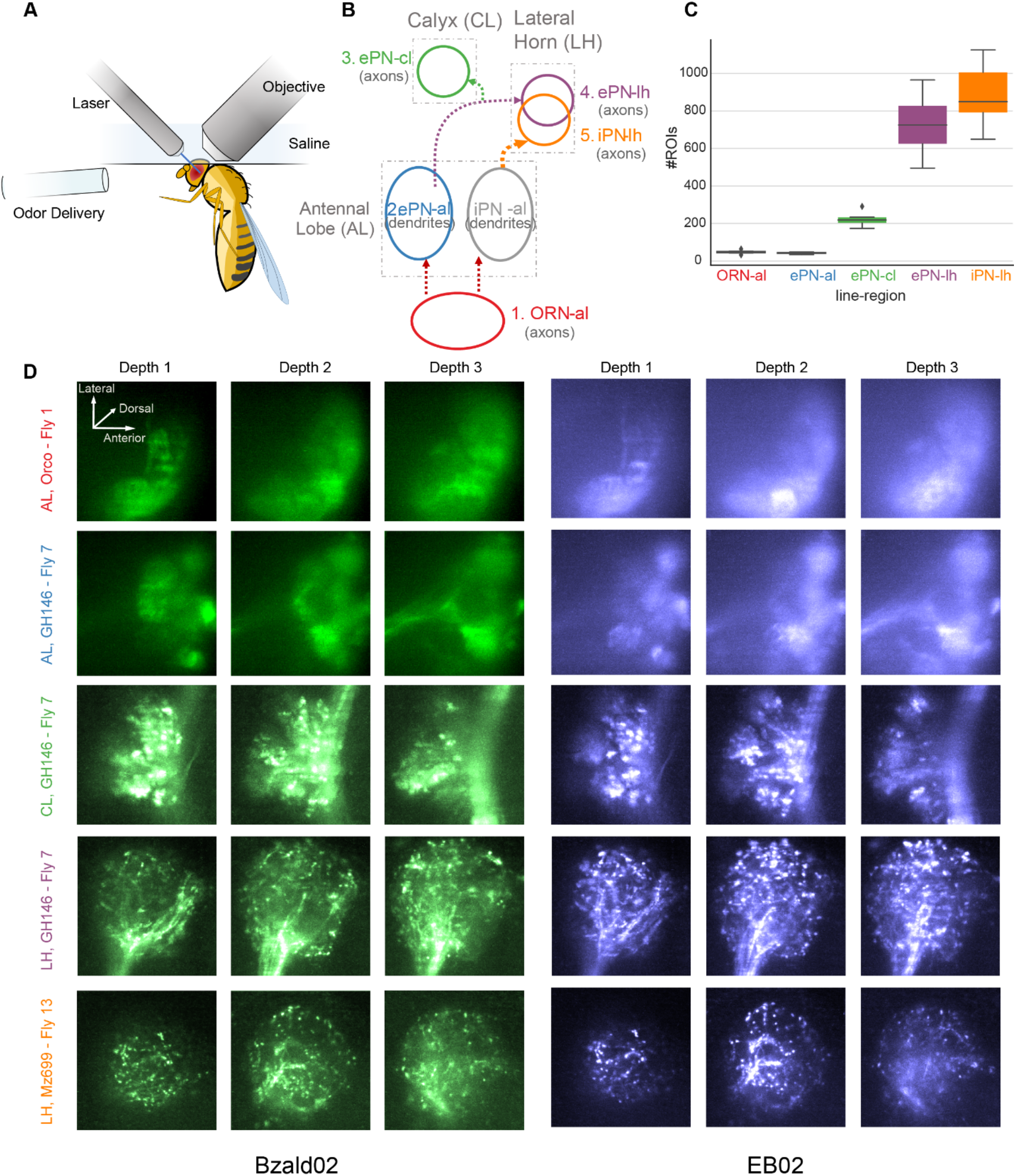
Light-sheet imaging for volumetric in vivo characterization of odor-evoked responses at the input and outputs of the antennal circuitry. **(A)** A schematic of the experimental setup. The fly is mounted on a custom mounting block with its antennae exposed to air stream and brain immersed in saline. At each scanning step, a whole brain plane is illuminated by a light-sheet with two wavelengths (488 nm and 561 nm). The fluorescent signals are collected by the objective and the downstream optical components. **(B)** Fly lines labeling any one of the following three distinct neural populations were used in our experiments: cholinergic ORNs expressing Orco co-receptor (ORNs), cholinergic projection neurons (ePNs) and GABAergic projection neurons (iPNs). For ORNs and iPNs, axonal activity alone was monitored. For ePN both dendritic responses in the antennal lobe and axonal responses transmitted onto mushroom body calyx and lateral horns were near simultaneously monitored. **(C)** The number of region of interest(ROI) extracted by a constrained non-negative matrix factorization algorithm is shown for different regions. Both the median and the interquartile ranges (IQR, 50%) are shown. Whisker lengths are 1.5 IQR past the low and high quartiles. Points out of this range were regarded as outliers. **(D)** Maximum responses observed during the Bzald0202 presentation window are shown for each optical plane. Each row shows changes in calcium activity from a labeled neural population at an anatomical location. Each column shows responses monitored at one depth of imaging stacks. **(E)** Similar plots as shown in panel D but now showing responses to EB02.

We probed the responses of ORNs, ePNs and iPNs to a panel of six odorants, each delivered at two concentrations. The odor panel was chosen to ensure diversity in functional groups, behavioral valence, activation patterns and concentrations (Badel et al., 2016; Knaden et al., 2012; Strutz et al., 2014). For example, benzaldehyde (*Bzald*) was reported to be repulsive (Ahsan et al., 2017; Strutz et al., 2014) and activate ventral glomeruli strongly compared with other stimuli(Badel et al., 2016), whereas ethyl acetate (*EA*) is regarded as an attractive cue that generates strong input to dorso-medial glomeruli (Ahsan et al., 2017). The light-sheet images acquired were segmented using an unsupervised non-negative matrix factorization method (Pnevmatikakis et al., 2016) (see **Methods** for details). Note that the ROIs corresponded to glomeruli for Orco-ORN axons and ePN dendrites (**Figure 2A;** top row), and ePN and iPN axonal boutons in calyx (CX) and lateral horn (LH) (**Figure 2A**; bottom row). A quick summary of the number of ROIs extracted from each fly is listed in **Figure 1C (** also refer **Supplementary Figures 1-5** for ROI masks that were extracted for each plane and in each fly).

**Figure 2:**
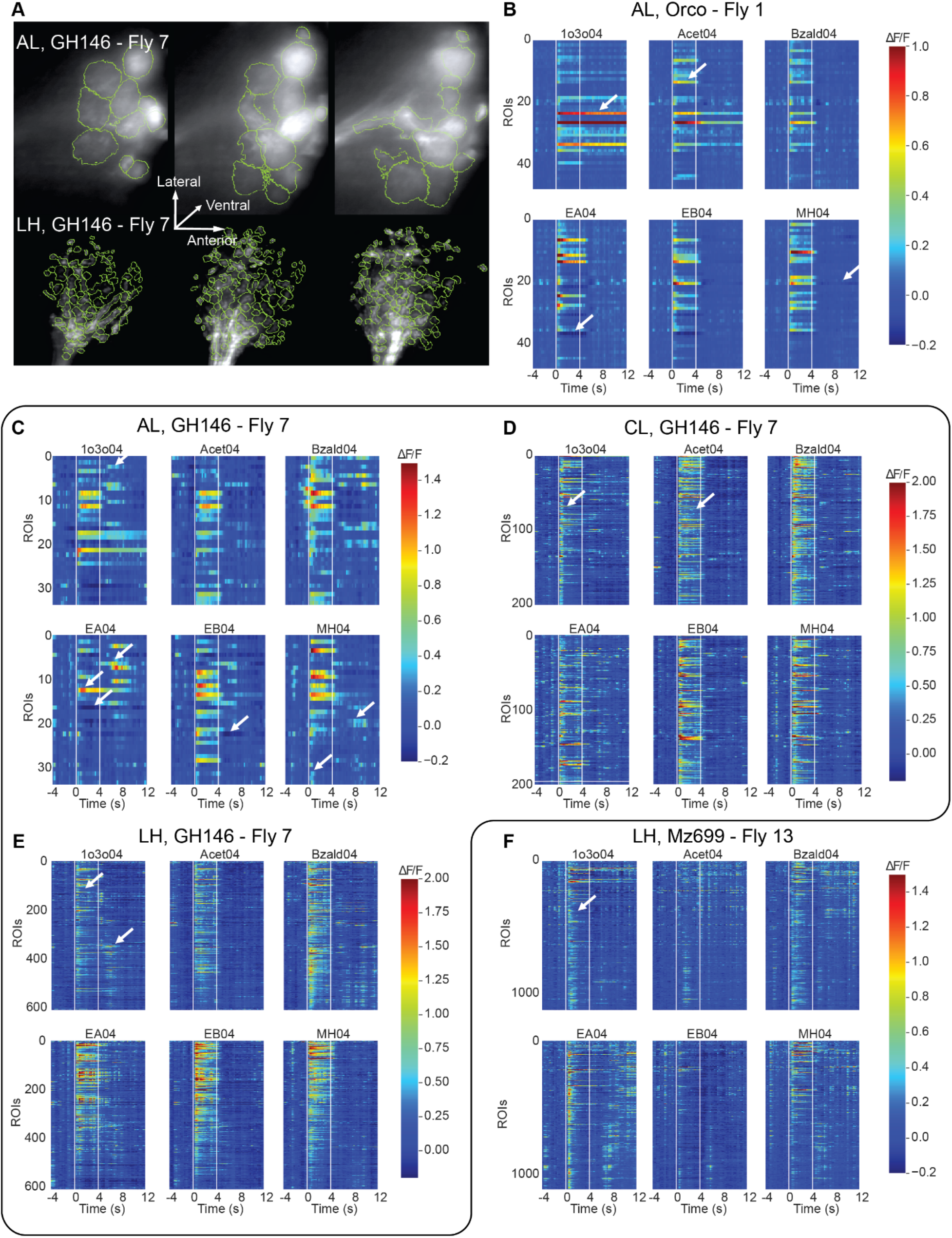
Extraction of spatial and temporal patterns of odor-evoked neural activity. **(A)** Region-of-interest (ROI) masks extracted by an unsupervised non-negative matrix factoriztion method are overlaid on top of raw calcium signals recorded from ePN dendrites in the antennal lobe (top panel) and ePN axons entering the lateral horn (bottom panel). Three panels are shown characterizing odor-evoked responses and ROI masks extracted at three different depths. Note the mask contours match the anatomical structures (glomeruli and axonal boutons) in both regions very well. **(B** through **F)** Representative responses to a panel of six odorants are shown as a data matrix. Calcium signals from individual ROIs extracted in each fly line/region are shown: olfactory receptor neurons in the antennal lobe (**B**); excitatory projection neuron dendrites in the antennal lobe (**C**); excitatory projection neurons axons in the mushroom body calyx (**D**); excitatory projection neuron axons in the lateral horn (**E**); inhibitory projection neuron axons in the lateral horn (**F**). Warmer color indicates stronger excitation, whereas cooler colors indicates inhibition. In each panel, each row represents temporal response of one ROI arranged in the order from dorsal to ventral. All the ROIs across different depths were pooled together and shown in the plot (from dorsal at the top to ventral planes at the bottom of each data matrix). Y-axis indicates the ROI numbers. White arrows annotate the typical response dynamics (see text for details).

In addition to large spatial coverage, we also acquired images rapidly (4 Hz sampling rate) to characterize odor-evoked, spatiotemporal response dynamics across the entire population of a specific type of olfactory neuron (**Figure 2**). Consistent with earlier reports (de Bruyne et al., 2001), we found that each odorant activated a unique combination of ORNs. For most ORNs, the sensory input lasted the duration of the odor response, and for certain odorant-ORN combinations, the unabated response persisted and outlasted the stimulus duration (**Figure 2B**; for example, *1o3ol04* and *Acet04*). In a few ORNs, substantial reduction in calcium signals were also evident during the odor presentation (ethyl acetate (*EA*) and ethyl butyrate (*EB*), **Supplementary Figure 6**). Prolonged excitatory responses, and inhibition that persisted after stimulus termination were pronounced at higher intensities (*MH02*; see **Supplementary Figure 7**). As the intensity was increased, additional ORNs were recruited for all odorants (**Supplementary Figure 7**).

In the downstream antennal lobe level, ePN dendrites showed richer response dynamics for all odorants (**Figure 2c**; **Supplementary Movie 1**). Increase in calcium signals after stimulus termination (i.e. ‘OFF responses’) were observed in many glomeruli. Consistent with prior results (Bhandawat et al., 2007), we also observed that odorants that evoked weak ORN inputs had amplified responses at the level of ePN dendrites (e.g. *Bzald04*). We also found the ePN signals attenuated more rapidly. More importantly, the mean response overlap between pairs of odorants appeared to remain consistent in all five fly-lines/regions examined (**Supplementary Figure 8**). Increasing odor intensity, recruited activity in additional glomeruli, and resulted in more complex changes in the response timing (**Supplementary Figure 7**).

As noted earlier, ePNs send axons to both the mushroom calyces and lateral horns, whereas iPNs project only to lateral horns. We found that activation patterns of ePN and iPN axons entering these higher centers were broadly distributed across several boutons. The responses tended to be more transient than even those observed at the level of ePN dendrites (**Supplementary Figure 6**). In most flies, the ordering of odorants based on strength of activation differed between the ePN dendritic and axonal compartments (**Supplementary Figure 9, 10**). Together, these results suggest that active signal transformation occurs between input and output compartments of these neurons. The activation became stronger for all odorants at higher intensity, but nevertheless remained highly transient and attenuated rapidly (**Supplementary Figure 7**). These observations remained consistent when data from across the flies were compared.

Note that these observations indicate that the light sheet imaging data remained consistent with several of the previous studies that only used a subset of the signals we recorded. However, our dataset allowed us to probe the spatial and temporal aspects of olfactory processing with greater resolution. Primarily, we sought to understand how sensory signals are represented and transformed at the input and output of the antennal lobe.

### Spatial organization of neural processes within the antennal lobe, calyx and lateral horn

How are functional units (ROIs) organized within each processing stage? Do ROIs that are spatially closer respond to odorants in a similar fashion? To examine this, we represented the response tuning of each ROI using a 12-dimensional vector, with each vector-component being the ROI’s mean response to an odorant (**Figure 3A**). Next, for every pair of ROIs, we computed response similarity (i.e. cosine of the angle between their 12-D tuning vectors) and plotted it as a function of spatial distance between them (i.e. distance between the two ROI’s centroids; **Figure 3A**). Note that a response similarity of 1 indicates that the two ROIs have very similar odor-evoked responses, whereas negative values indicate response tunings that are opposite.

**Figure 3:**
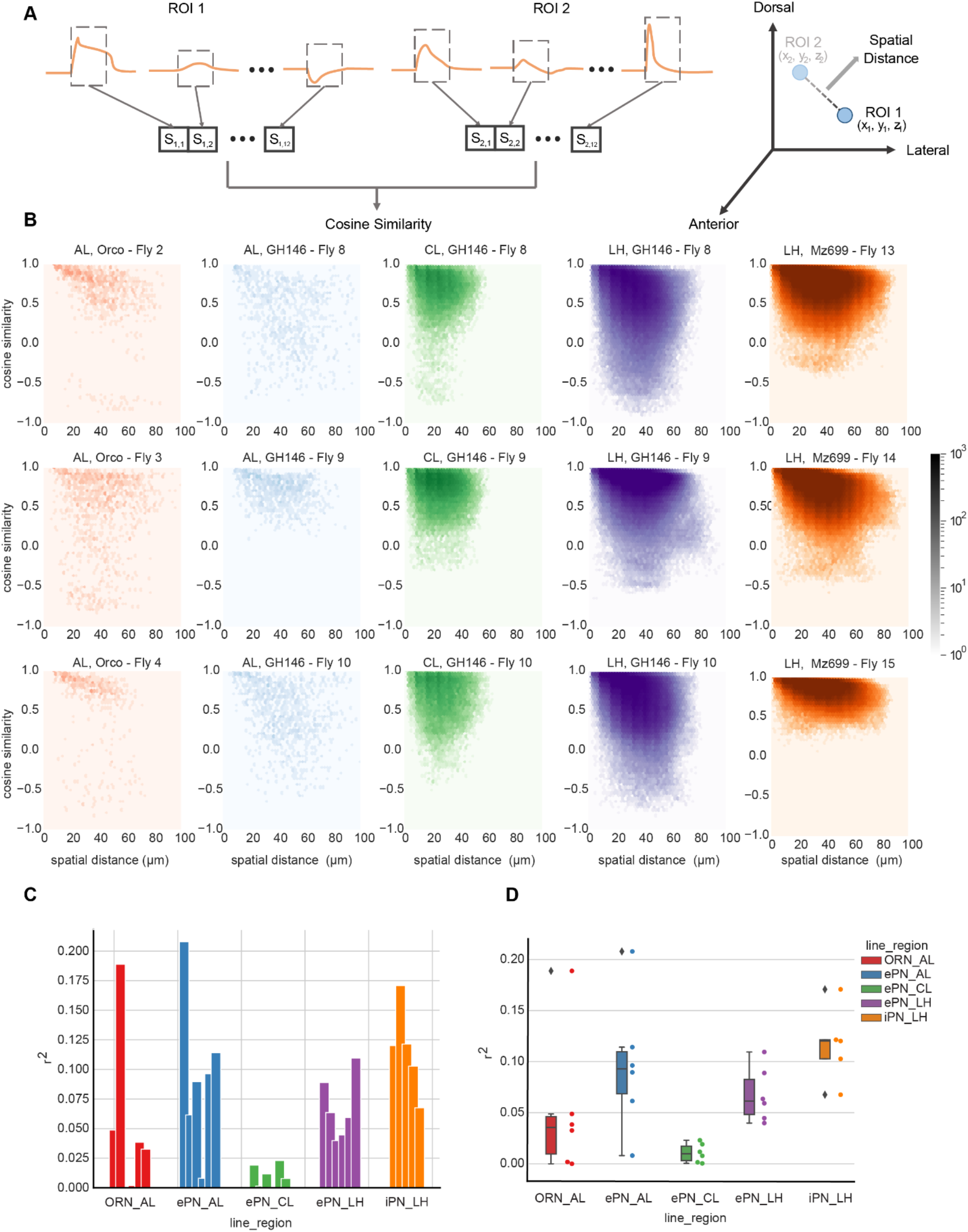
Functional distance vs. spatial distance. **A** schematic illustrating how functional distances (left) and spatial distances right) were calculated. For each ROI, its tuning vector consists of twelve elements. Each vector component represents its mean response (over time) to one odor stimulus. Functional distance was calculated as the cosine similarity between two ROI’s response vectors. Spatial distance between a pair of ROIs was calculated as the Euclidean distance between two ROIs spatial location as shown in the right panel. **(B)** A scatter plot showing relationship between functional distance (y-axis) vs. spatial distance (x-axis) for all ROI pairs. Each column indicates an anatomical region. Results from three representative flies are show for each line (three rows). For all three lines, there was a weak but general trend that spatially near-by ROIs have higher correlation between their odor-tuning vectors. **(C)** The linearity of functional vs spatial distance relationship was quantified (coefficient of determination or r^2^) and shown. Each bar indicates the r^2^ value of a linear regression model, with spatial distance as independent variable and functional distance as dependent variable for one region and from one fly. Colors correspond to different regions matching the color scheme shown in **panel b**). **(D)** Same as **panel c**, but coefficient of determination summarized as box plots.

Our results indicate that for all three neural populations examined (ORNs, ePNs and iPNs), there was a weak but general trend that spatially near-by ROIs were similar in their odor tuning (**Figure 3B**). Notably, the ‘space vs. tuning’ distributions were different between ORN axons and ePN dendrites in the antennal lobe, and between ePN and iPN axons innervating the lateral horn. The former result indicates active transformation of sensory signals as it propagates through the glomerular microcircuits in the antennal lobe, while the later observation suggests that ePN and iPN innervations in the lateral horn follow different organization principles. However, in all flies examined, there was barely linear spatial organization in the calyx (**Figure 3B** 3^rd^ column).

Taken together, these results indicate that, similar to results in the mouse olfactory bulb (Ma et al., 2012), the spatial organization of odor representation in fly antennal lobe is weak. Notably, this organizational feature was present in all flies examined (**Figure 3C-D**).

### Characterizing spatial organization of odor tuning across neural populations and across flies

To better understand the spatial organization of ROIs in different regions, we positioned each ROI based on its XYZ co-ordinates in the fly brain and colored it based on its odor response tuning. As mentioned earlier, the specificity or tuning of each ROI was defined using a 12-D vector (each vector component to indicate the response elicited by each of the twelve stimuli used; **Figure 4A**). The 12-D ROI tuning vectors were dimensionality reduced to a 3D space using multiscale scaling (MDS) algorithm (**Figure 4a**). Each 3-D MDS vector was then assigned a color using a 3-D RGB color scale (see **Methods**). Note that ROI’s with similar tuning profiles were assigned similar colors. Furthermore, to create suitable points of reference or ‘tuning landmarks’, a few artificial templates were generated, and the colors each one of these templates was assigned is also shown (**Figure 4B; Supplementary. Figure 11** shows colors that were assigned to a more elaborate set of reference vectors/templates).

**Figure 4:**
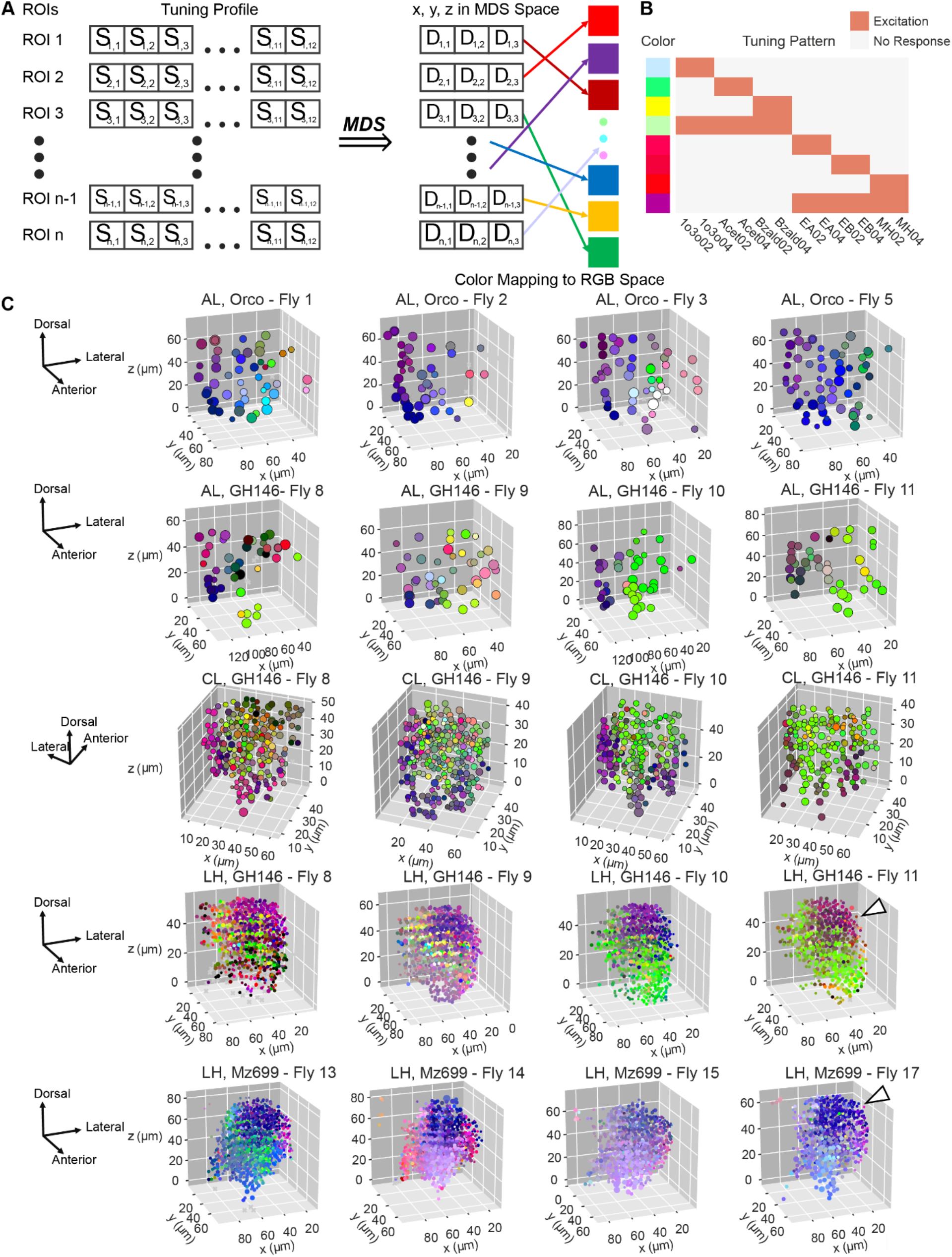
Spatial organization of extracted ROIs. **(A)** An ROI’s tuning profile is again summarized as a 12-dimension vector as in **Figure 3a**. To obtain the 3D color space, we used multidimensional scaling (MDS) to map the pairwise 12-D functional distance in the tuning space onto a 3D color space. Then the color of each ROI was assigned based on the coordinates in the 3D color space where each axis corresponded to red/green/blue colors (**Methods**). **(B)** The correspondence between tuning profiles and colors are shown. Color bar on the left indicates the color correspondence of the ‘artificial landmark’ tuning profiles shown on the right (same row). Green/Yellow colors indicate ROIs that are more responsive to aversive odorants (1o3o, Acet and Bzald). On the contrary, Red/magenta colors show ROIs tuning preference to attractive odors (EA, EB and MH). **(C)** ROIs are shown in their actual 3D spatial locations across different regions/fly lines (4 representative flies are shown for each region). Each ROI is also labeled by the color obtained from MDS analysis indicating tuning properties.

Using this odor tuning-based coloring approach, we visualized each ROI and compared their stimulus specificities (**Figure 4C**). Note that red through dark purple/dark blue colors identify ROIs that were strongly activated by attractive odorants (*EA04*, *EB04* and *MH04*; **Figure 4B**), where the green through yellow colors identify ROIs that responded to the repulsive ones (*Bzald04*, *1o3o04*, *Acet04*; **Figure 4B**).

Consistent with prior studies, in *orco*-labelled flies (**Figure 4C**; first row), we found that dorso-medial and ventro-medial glomeruli were activated strongly by attractive odorants (*EA04*, *EB04* and *MH04*; blue/purple colored ROIs). Whereas glomeruli in ventro-lateral regions tended to respond more to the repulsive odorants (green ROIs). It is worth noting that the attractive odorants evoked strong responses and activated more glomeruli at the level of sensory neuron axons.

In comparison, at the level of ePN dendrites (**Figure 4C**; second row), the odor tuning maps changed. First, the extent of activation of the attractive odorants was restricted to fewer glomeruli located in the dorso-medial and ventro-medial regions. Response to the repulsive odorants, that were weaker at the level of Orco sensory neuron axons, became stronger and spread to more glomeruli in the ventro-lateral regions (note that the response amplification to *Bzald*, *Acet* and *1o3o* is also evident in ePN PSTH’s shown in **Supplementary Figure 9, 10**).

In the calyx, we found that the ROIs in the core region differed in tuning from the ROIs that bordered them and formed the outer-rim. Attractive odorants strongly activated the outer-rim, whereas ePN axons entering the core strongly responded to repulsive odorants. These results are again consistent with pure anatomical studies that have examined how a few glomeruli in the dorso-medial region of the antennal lobe innervate the calyx (Tanaka et al., 2004).

Finally, in the lateral horn too we found that both ePN and iPN axons were spatially organized based on their odor tuning. While all repulsive odorants evoked strong responses at the level of ePN axons in the lateral horns, iPN axons only weakly responded to some of those odorants (for example iPN axonal responses to *Acet04* were weaker in all flies; refer **Supplementary Figure 9, 10**). Intriguingly, a stereotyped region in the dorso-lateral lateral horn received ePN and iPN axons that matched in their response selectivity (blue-purple region indicating response to attractive odorants). While in the rest of the lateral horn, the ePN and iPN differed in their response tuning. This suggests that matched feed-forward excitation and inhibition may compete in the lateral horn regions receiving inputs regarding attractive odorants, while interactions between mismatched excitatory and inhibitory inputs may occur in other regions.

### Relating dendritic ePN inputs with their axonal outputs to higher centers

Next, we investigated the relationship between the ePN responses in the antennal lobe (dendrites) and those transmitted to mushroom body calyx ad lateral horn (axons). Previous connectomic studies had shown that each ePN project its axonal terminals to a limited number of locations in the calyx and lateral horn (Zheng et al., 2018). This wiring pattern would suggest that each ePN may simply send its output to a spatially restricted region downstream, and may have only a minimal influence on functional signals reaching the other spatial loci. Since we acquired data from ePN dendrites and axons near simultaneously from each fly, we examined if this was indeed the case.

We performed a regression analysis to understand the functional relationship between ePN input and output compartments (see **Methods; Figure 5A**). In this approach, we used linear combination of ePN dendritic responses to predict the responses at each individual axonal bouton. Note that each row of the regression weights matrix (**Figure 5B**) indicates how the regression weights from multiple antennal lobe ROIs were linearly combined to map onto each calyx or lateral horn ROI. Alternately, each column of the weight matrix show in **Figure 5B**, can be interpreted as the contribution each antennal lobe ROI makes in generating axonal responses in the two downstream regions.

**Figure 5:**
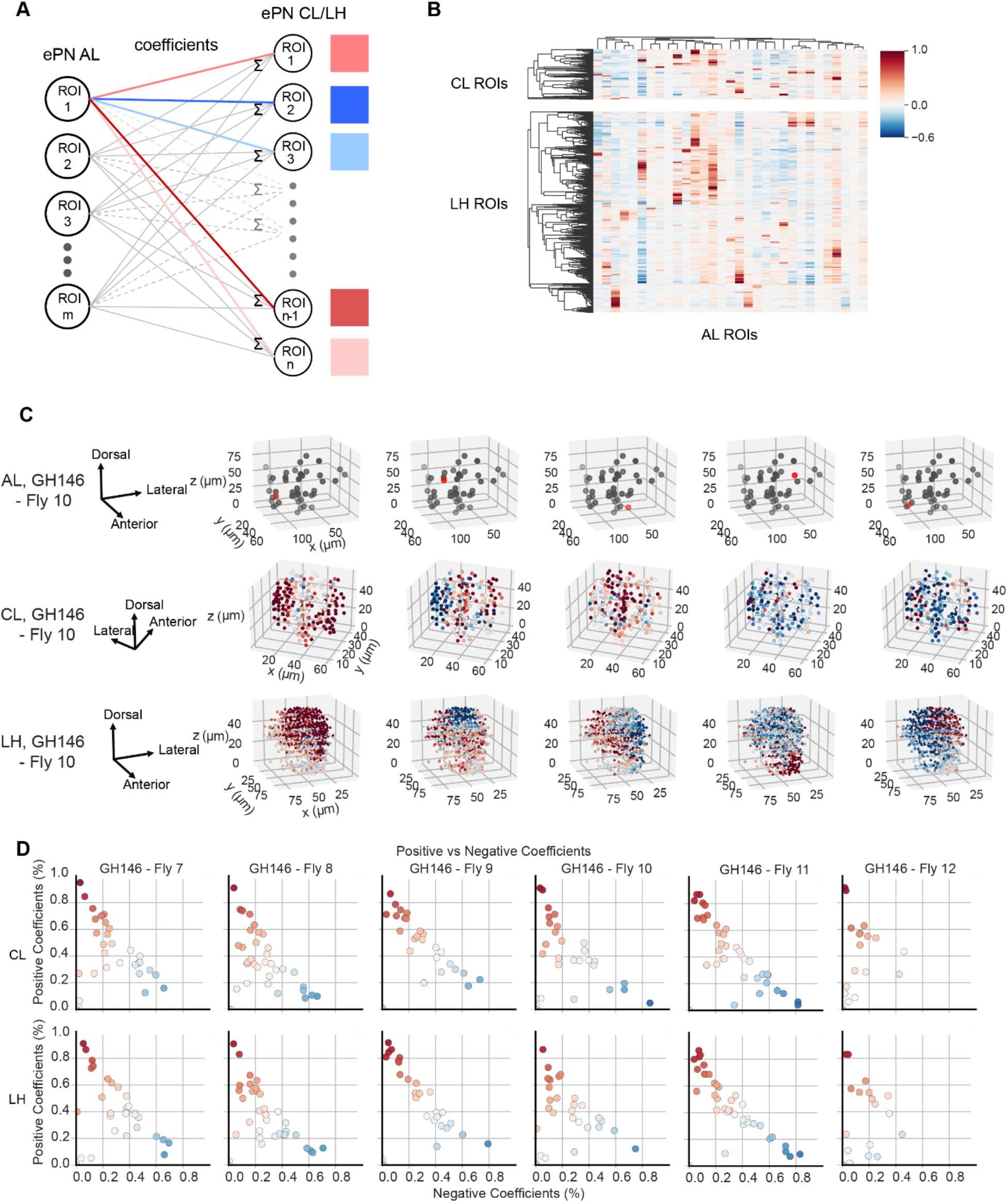
Linking dendritic inputs of ePNs with their axonal outputs (I/O mapping). **(A)** The schematic shows how linear regression was performed to obtain the coefficients relating responses in two regions (input – antennal lobe; output – calyx/lateral horn). Responses over time of each axonal bouton/ROI in the lateral horn/calyx regions were predicted using a linear combination of ePN dendritic responses. Regression weights were learned using a multi-task lasso regression (see methods). **(B)** The regression coefficients learned from a representative fly are shown. Each column corresponding one ROI in AL as regressors. Each row shows the weights that were assigned to different ePN dendritic ROIs used to predict response a single ROI in the LH/CL. Only non-zero columns are shown here. Warmer color indicates stronger positive influences and cooler color shows stronger negative influences. **(C)** 3D scatter plots showing single antennal lobe ROI’s functional influence on the ePN axonal responses observed in the calyx and lateral horn. The first row shows the spatial location of the specific ROIs in the antennal lobe (ROI labeled in red). Rows two (calyx) and three (lateral horn) show ROIs in these locations colored using the regression coefficients obtained. Each column identifies the AL ROI (first row) and its influence in calyx and lateral horn (rows two and three, respectively). The orientations of each imaged region are indicated on the left panel. **(D)** The percentage of significant positive (Y-axis) and negative coefficients (X-axis) assigned for each antennal lobe ROI are plotted against each other. Color encodes the net difference between the positive/negative coefficient percentages; for instance, warmer colors represent that the antennal lobe ROI had more positive coefficients than the negative ones. Results from 6 experiments/flies are shown in different columns. Top row shows regression weight distribution in the mushroom body calyx, and the bottom row reveals similar results but in the lateral horn.

Contrary to our expectations, each ePN dendritic activity showed a more global contribution downstream (**Figure 5C, D**). As can be noted, most columns have both hot (positive influence) and cool (negative influence) colored vector components indicating that majority of antennal lobe ROIs had a mixed influence in calyx and lateral horns axonal responses (i.e. positive influence in some regions and negative influence in others). Only a few antennal lobe ROIs had predominantly positive or negative influences on the downstream regions. Note that, for some antennal lobe ROIs, the ratio of positive to negative influence also varied between calyx and lateral horn (**Figure 5C;** 3^rd^ column). This observation implies that the input from the antennal lobe is restructured differently between the two downstream targets.

To understand the spatial distribution of how each antennal lobe ROI contributed to downstream activity, we mapped the vector of regression weights onto the spatial locations of each axonal bouton (**Figure 5C**). The antennal lobe ROIs had diverse functional relationships with the ePN axonal responses observed in the calyx and lateral horn. Nevertheless, the regression weights from a single antennal lobe ROI appeared to be spatially organized, with regions of positive and negative influences occurring in spatially contiguous regions juxtaposed next to each other. This spatial arrangement was much clearer in the lateral horns and to a lesser extent also observed in the calyx. Interestingly, antennal lobe ROIs that were spatially close to one another had functional innervation patterns that were markedly different from one another (**Figure 5C**; columns 1 vs column 5 shows functional mapping of inputs from two ROIs in the dorso-medial antennal lobe)

To quantitatively compare the influence different antennal lobe ROIs had on the two downstream regions, for each ROI, we plotted the fraction of positive influence/weights versus the fraction of negative influence/weights (see Methods; **Figure 5D**). Note that ROIs that were close to the two axes had predominantly either positive (closer to y-axis) or negative (closer to the x-axis) influence. Most ROIs had a mixed influence and were positioned away from both these individual axes in these plots. Notably, a similar distribution of ePN antennal lobe ROI weights were observed in both calyx and lateral horns, and across different flies. In sum these results indicate that the functional relationships between responses observed in the dendritic and axonal ePN compartments are complex, and diverse.

### Temporal patterning of odor-evoked responses

So far, we have examined how odor-evoked responses are spatially distributed at the level of ORN axons and how these responses map onto the two downstream neural populations (ePNs and iPNs). Next, we sought to examine how these odor-evoked responses are patterned over time. Our results indicate that spatial patterns of activity in the antennal lobe, both at the level of *Orco* axons (**Supplementary Figure 12**) and ePN dendrites (**Figure 6A**), were highly similar immediately after the onset of the odorants. However, over time these spatial patterns of neural activity evolved to become more distinct.

**Figure 6:**
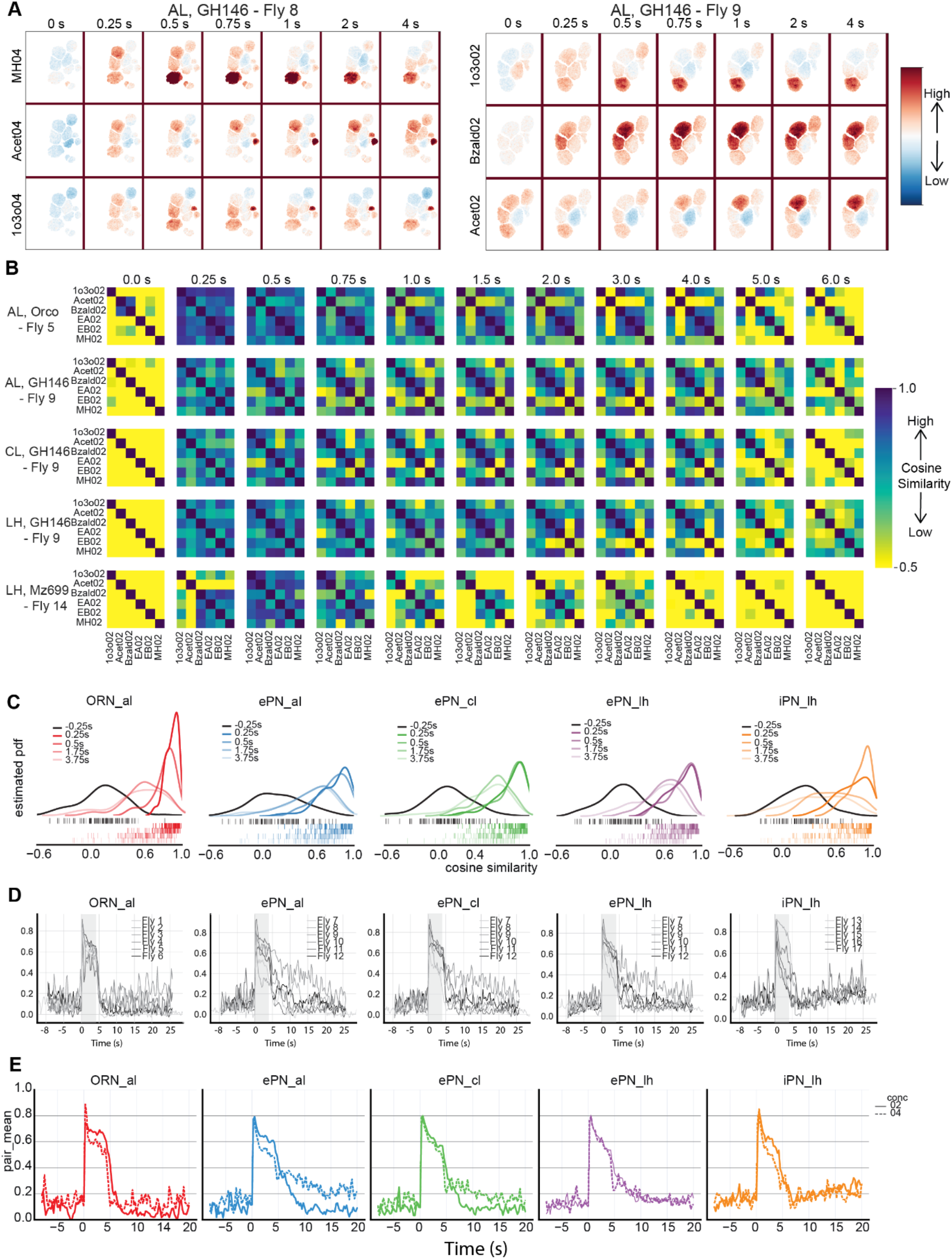
Odor-evoked responses decorrelate over time. **(A)** Change in fluorescence signals (ΔF/F) for a few representative ROIs on single optical plane in the antennal lobe are shown as a function of time since odor onset (shown at the top of the panel). Each row reveals responses evoked by an odorant. Right panel show evolution of odor-evoked responses in the antennal lobe ePN dendrites observed in another fly. **(B)** Pattern similarity matrices for a representative fly for each labeled fly line/region are shown. Each element in the matrix is the cosine similarity value between a pair of odorants. Hot colors indicate stronger similarity, and cooler colors indicate weaker similarity. Each row reveals how pairwise odor similarities evolve over time. Again, time since odor onset is indicated at the top of the panel. In total, pairwise similarity matrices at eleven time points are shown. Odor stimulus was presented from 0.0 sec to 4.0 sec. Note that similarity matrices with higher pattern similarities (cooler/blue colors) at the start of response and gradually decorrelate over time (hotter/yellow colors). This can be observed in all five rows corresponding to responses observed in ORNs, ePN dendrites, ePN axons in the calyx, ePN axons in the lateral horn and iPN axons in the lateral horn. **(C)** Distributions of pairwise pattern similarity (cosine distance) obtained using kernel density estimation (see **Methods**) are shown. Each curve shows pairwise pattern similarity distribution at one time point. In each panel, response similarity distributions are shown for five different time points before and during stimulus presentation. Tick marks shown below the distributions represent pairwise similarity between every pair of odorants and across flies. Ticks are color coded following the same scheme used for the distributions shown on the top. **(D)** Mean pair-wise cosine similarity in each region is shown as a function of time. Each trace shows the mean cosine similarity value across all odor pairs for each individual fly. Color bar indicates the 4 s duration when the odorant was presented. Five panels are shown to illustrate results from the three fly lines used in the study. **(E)** Mean pair-wise cosine similarity as a function of time is shown. Two traces, corresponding to the two concentrations of odorants used, are shown tracking changes in mean cosine similarity across odorants/flies.

To quantify this observation, we computed the cosine similarity between responses evoked by different odorants at specific time point during stimulus presentation (**Figure 6B**; see **Methods**). As can be observed, the responses evoked by different odorants at all five neural processes (Orco axons, ePN dendrites, ePN axons entering calyx and lateral horn, iPN axons entering lateral horn) had high correlation immediately after the onset of stimulus. However, over time these correlations reduced and responses evoked by different odorants became more distinct from each other (i.e. lower correlations/similarity).

These observations were further corroborated when pairwise similarities between odorants across flies were examined (**Figure 6C**). Note that pairwise similarities between most odorants immediately after onset were high in all three lines examined (tick marks shown below the probability density functions in **Figure 6C**). The pre-stimulus activity before onset of any two stimuli showed wide dispersion of cosine similarity values with a mode near zero indicating randomness in signals recorded during this time period. Immediate after odor onset, the distributed shifted right indicating an increase in odor similarity across pairs of odorants and observed in all flies examined. With progression of time, the distribution of pairwise cosine similarities shifted leftwards (i.e. towards lower values) indicating decorrelation of odor-evoked responses.

The evolution of mean pair-wise correlation across odorants over time showed variable reduction rates in each individual fly examined (**Figure 6D**). As can be expected, in all three neural populations, low concentration stimuli decorrelated faster and more than responses to the same set of stimuli evoked at a higher concentration (**Figure 6E**). Interestingly, only in the ePN axonal projections the speed of response decorrelation was comparable at both low and high concentrations. This result directly suggests that some additional modification of response patterns occurred in this neural population to rapidly make the neural activity evoked by each odor more distinct from others (**Figure 6E**).

Taken together, these results indicate that the odor-evoked response patterns and the discriminatory information needed for selective recognition evolve over time in the early fly olfactory circuits. Consistent with findings from other model systems (Friedrich and Laurent, 2001; Gschwend et al., 2015; Raman et al., 2010), the observed temporal patterning made odor-evoked response patterns to become different from the initial stimulus-evoked activity but also more distinct when compare to other odorants.

### Idiosyncratic processing underlies how odorants are segregated over time

Given that the initial olfactory circuits have been reported to be stereotyped across flies (Fishilevich and Vosshall, 2005; Jefferis et al., 2007; Vosshall et al., 2000; Vosshall, 2008), it would be reasonable to expect the variability across flies in these peripheral neural circuits to be low. However, our results (**Figure 6D**) indicate that decorrelation of odor-evoked responses occur at different rates in different flies. To further examine this issue, we compared how similarity between pairs of odorants evolved over time in different flies (**Figure 7A**). Note that the hot colors indicate high correlations/similarity and cool colors indicate negative correlations. Also, clearly observable in the correlation plots shown for the two representative flies is the initial vertical band of high correlation immediately after odor onset. However, note that correlation between different stimulus pairs transformed rapidly. Bands of highly-correlated responses observed immediately after odor onset (show using hotter colors) transitioned to dissimilar responses (less hot colors) at varying points in time. More importantly, the pairwise odor correlation patterns differed between flies indicating that although the odor responses became more distinct, which pairs of odorants became separable at which point in time depended not only on the odorants but also varied from one fly to another.

**Figure 7:**
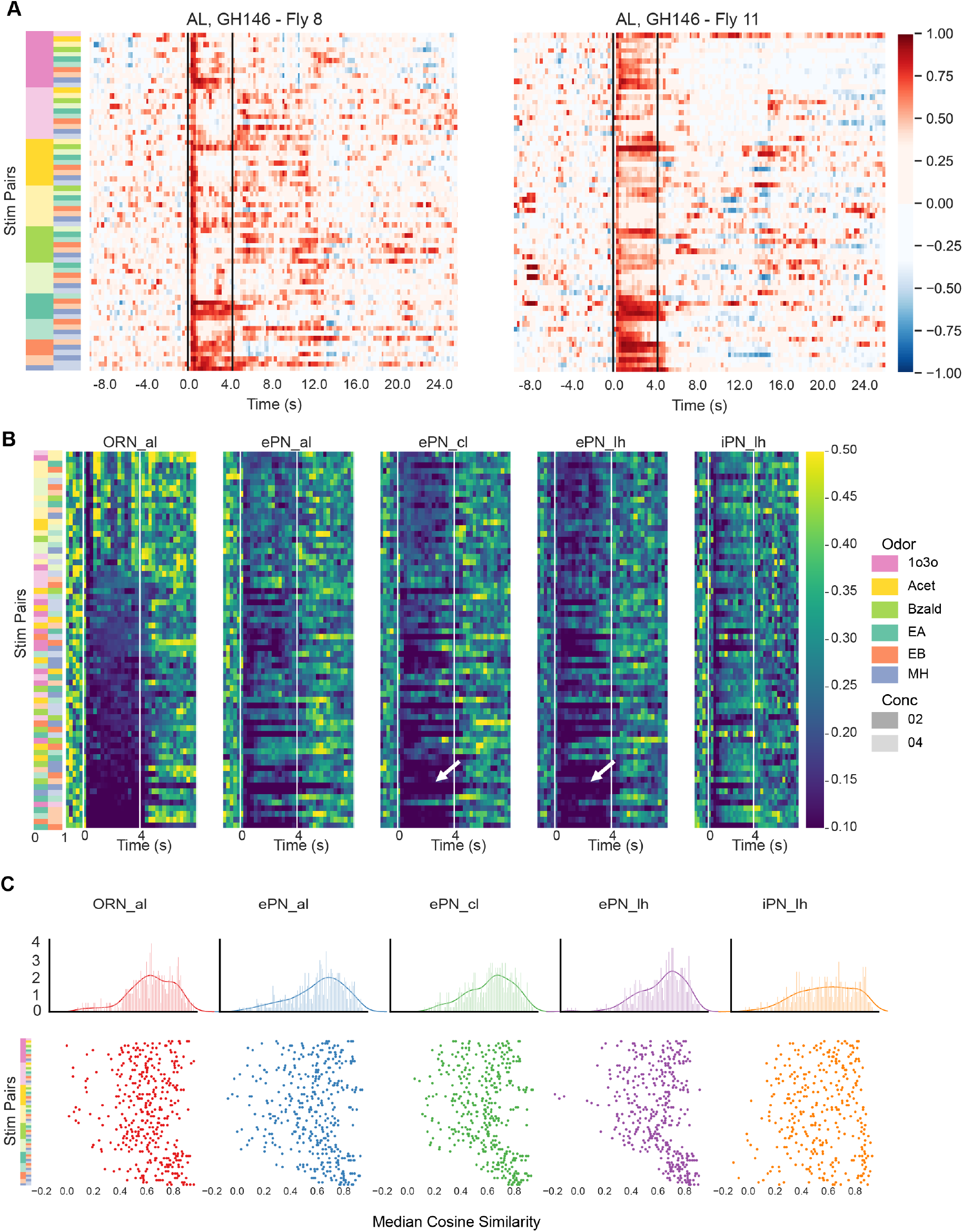
Pairwise odor similarities vary across flies. **(A)** Pairwise cosine similarities of ePN dendritic responses and how they evolve as a function of time are shown as a heatmap. Each row tracks response similarity between one odor pair, and each column represents one time point. The identity of each stimulus pair tracked in a given row is indicated using a color bar on the left of the heatmap. The four second window when the odorant was presented in indicated using black vertical lines. Hotter colors indicate more similarity and cooler colors indicate less similarity. Panel of the right, shows evolution of pairwise cosine similarity for the same pairs of odorants (ordered as shown on the left panel) but in a different fly. **(B)** For each odor-pair, the standard deviation in pairwise odor similarity across individuals were calculated and plotted as a function of time. Hot regions in the heatmap show the standard deviation of the cosine similarity across individual fly was greater (i.e. more variability across flies). Similar plots, but characterizing variation in pairwise odor similarity in the five fly line/regions studied are shown. The color bar on the left identifies the odor pair tracked in each row. Note that the rows are sorted in descending order based on standard deviation values observed in the ORN level. **(C)** The median cosine similarity during the 4s stimulation period for each stimulus pair, and for each fly, is shown as a scatter plot (bottom). Therefore, each marker represents median pairwise odor similarity observed in a single fly, and each row tracks variation across flies. The identity of the odor pairs corresponding to each row is indicated using the color bar on the left. Tighter packing of individual markers along a single row indicates responses observed across individual flies were highly reliable. The overall distribution across odor pairs and flies is shown on the top.

To further quantify this result, we computed and plotted the standard deviation in pairwise odor response correlations across flies (**Figure 7B**). High standard deviation would identify pairs of odorants that were decorrelated differently in different flies. Our results indicate that some odor pairs were indeed processed in a relatively conserved manner across flies (identified using arrowheads), whereas many differed starting from the activity they evoked at the level of ORN axons. The standard deviation between flies were relatively less at the level of ePN axons compared to their dendritic activity, whereas the multiglomerular iPNs had the higher levels of variability even though they integrated inputs from multiple different ORNs. These results indicate that while odor-evoked response patterns decorrelated to become more distinct over time in all flies, this computation was performed in an idiosyncratic fashion.

To illustrate the variability across flies, for each stimulus pair we plotted the median response similarity (**Figure 7C;** median over time and each row shows variance across flies for each odor-pair). Our results indicate that the attractive odorants (indicated using arrowheads at the bottom of the panel) were more reliably represented across flies and evoked less variable responses in ORNs and ePNs. Overall, the variability was reduced at the level ePN axonal responses in calyx and lateral horns. In sum, these results indicate that odor-evoked responses, even in the early olfactory circuits are not stereotyped for most odorants.

### Stimulus evoked ON and OFF responses

Finally, we examined how stimulus-evoked responses were patterned after the stimulus termination (i.e. the stimulus-evoked OFF responses). We found that at the level of ORNs two types of responses were observed after stimulus termination: continuation of the ON response and inhibition in new ROIs that did not have an ON response. Excitatory responses only during the OFF period were seldom observed at the level of sensory neuron responses (**Figure 8A**).

**Figure 8:**
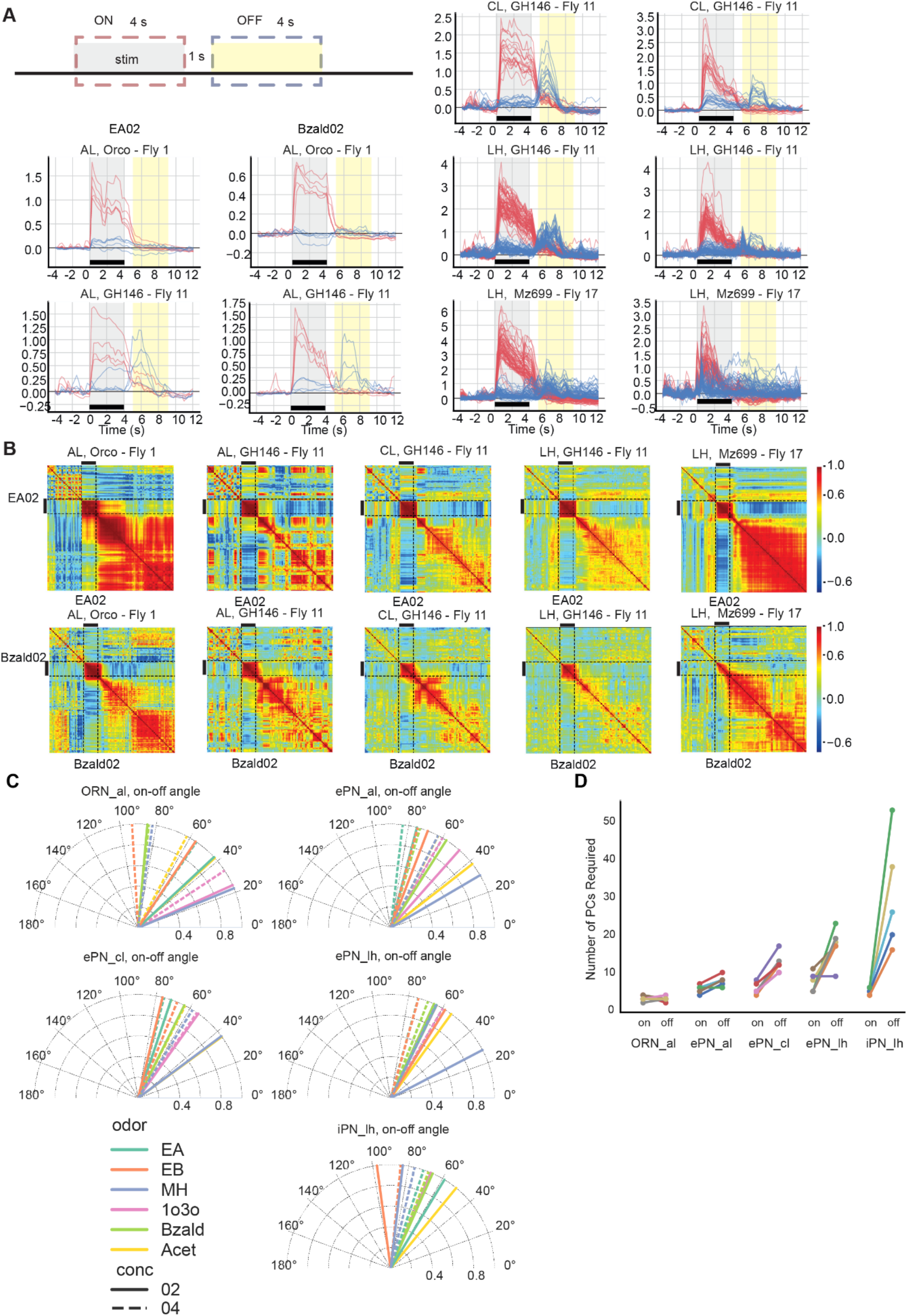
Odor evoked ON vs. OFF responses. **(A) The** top and bottom 5% of traces sorted by the mean amplitude during stimulus are shown, with the top 5% in red, and the bottom 5% in blue. The ON and OFF response windows are schematic schematically identified in the plot. Responses evoked by two representative odorants in each of the five fly line/region combinations are shown. **(B)** Evolution of correlation between neural activity before, during and after odor exposure are shown as a heatmap. The black bar on the left and top indicates the time period when the stimulus was delivered. Hot colors indicate high similarity and cool colors indicate low similarity. Note that each non-diagonal pixel represents similarity between ensemble ROI activities in one time bin versus those in another time bin. One row or column represents the correlation between one ensemble ROI activity vector with all other ensemble ROI vectors. Correlation heatmaps for two representative stimuli are shown for all three fly lines and five locations imaged. **(C)** Angle between mean ON and OFF response patterns evoked by each odorant is shown. Different colors represent different stimuli and the line style represents the two concentration levels. **(D)** The number of principal components needed to account for 90% of the data variance during ON and OFF response periods are plotted as a pair of points for each fly line/regions. Colors indicate individual flies.

In comparison, the OFF responses observed at the level of ePN dendrites and ePN/iPN axons showed response patterns that were more orthogonal with respect to the ON responses (**Figure 8A; Supplementary Figure 13**). ROIs that were active during ON period returned to baseline activity levels or even below baseline level responses (i.e. inhibition) in many ROIs. Whereas, ROIs that were not activated by stimulus exposure or even inhibited during the ON periods, tended to have a strong OFF response.

To understand how dissimilar were the neural responses observed during and after stimulus termination, we performed a cross-correlation analysis. A snapshot of activity across all ROIs was regarded as a high-dimensional vector. The similarity between each response vector with every other response vector that was observed over time was computed and shown succinctly as a correlation matrix (**Figure 8B**). Hot colors were used to indicate high correlation/similarity and cool colors to indicate negative correlation/dissimilarity. Note that while response vector observed during odor presentations (i.e. the ON responses) were well correlated amongst themselves, and the responses observed after odor termination (i.e. the OFF responses) poorly correlated with these ON responses (arrow head). This relationship between the ON and the OFF responses was observed in all three neural populations and in every fly studied.

To quantify how much the OFF patterns deviated from the ON patterns, we computed the angles between the mean population vectors during the ON and OFF periods (**Figure 8C**). Consistent with interpretation of the correlation plots, for most odorants, the ON and OFF response vectors evoked by the same odorant had an angular similarity in the 60°– 100°range (closer to 0° indicates similar responses and 90° indicates orthogonal responses).

Finally, we examined whether the response patterns evoked after odor termination are as diverse as those observed during stimulus presence. To compare pattern diversity, we used the number of principal components that were required to capture 90% of the total variance of the data (can also be thought of as a measure of intrinsic dimensionality of the dataset; **Figure 8D**). Surprisingly, compared to the ON responses, our results indicate that the OFF patterns were more diverse and needed more principal component to capture the same amount of variance in the response patterns observed.

In sum, our results indicate that for most odorants, another round of diverse response patterns were observed following stimulus termination. More importantly, these response patterns were dissimilar to the odor-evoked ON responses, and were a common encoding feature in all three neural response populations and all flies studied.

## Discussion

We sought to understand how sensory input from olfactory receptor neurons are spatially and temporally reformatted by two different downstream neural populations: ePNs and iPNs. While ePNs are cholinergic and receive input from a single glomerulus (Couto et al., 2005), iPNs are mostly GABAergic and multiglomerular (Wang et al., 2014). Further, while ePNs project to both calyx and lateral horn, iPN axons only innervate the lateral horns (Strutz et al., 2014). So, given the differences in the nature of input received (from one vs. many types of ORNs), and the downstream centers they feed onto, it is reasonable to expect that the ePNs and iPNs use different transformations to reformat sensory information received. However, our data reveal that several spatial and temporal aspects of odor-evoked responses were strikingly similar in both these neural populations.

### Spatial Organization of ePN and iPN processes

Our results indicate that both ePN and iPN axons were organized in the lateral horns such that nearby spatial regions had similar odor tuning. Though this relationship was weak, it was still significantly higher than the spatial organization of ePN axons in the calyx. More importantly, our results indicated that in lateral horn, ePNs and iPNs axons with similar stimulus tuning spatially overlapped. Since the iPN axons in different regions of lateral horn were differentially tuned to different odorants, our results indicate that this neural population may provide feed-forward inhibition in an odor specific manner.

In the antennal lobe, the ePN dendrites again showed a weak correlation between odor tuning and spatial location. Notably, the tuning vs distance relationship varied between flies. The weak spatial organization of antennal lobe neural activity in flies are qualitatively similar to results reported in the mice olfactory bulb (Ma et al., 2012). In the calyx, the ePN axons were organized such that the attractive odorants strongly activated the periphery, whereas the repulsive odorants were driving responses in the core regions. This organizational structure was found in all the flies, and is consistent with anatomical studies that revealed that dorso-medial glomeruli innervate the outer rim of the calyx and the ventral glomeruli send processes to the inner core regions (Tanaka et al., 2004). Note that this organization of ePN axons in the calyx is indeed non-linear and therefore was not picked up in the linear correlation measures we used to quantify the relationship between ROI location and tuning. Taken together, these results indicate that the observed differences in the organizational logic between dendritic and axonal compartments of ePNs was observed in all flies examined, arguably may indicate that different computations (global vs. local) that may be performed in these centers.

We also examined whether a single antennal lobe region had a one-to-one, local or global influence on the downstream centers. Note that the activity observed in the axonal boutons entering calyx and lateral horns incorporates the feedforward input from the antennal lobe ePNs and any recurrent pre-synaptic inhibition that is recruited in the target region. Although our results were obtained from a linear statistical analysis with a sparsity constraint, it indicates that each antennal lobe ROI contributes globally. Furthermore, most ROIs appeared to have both positive and negative influence in the downstream regions indicating that ePN activity is further transformed as it reaches calyx and lateral horns. These results were again replicated in different flies indicating that this is a generic organizing principle in the fly olfactory system.

In the lateral horn, a sterotyped, dorso-lateral region that was activated by all putative attractive odorants were detected. A prior study had identified a similar region in the lateral horn for the iPN axons (Strutz et al., 2014). Our results reveal that this lateral horn region is not only innervated by feed-forward inhibition (i.e. iPN axons), but also by feed-forward excitatory inputs (i.e. ePN axons) from the antennal lobe as well. Such overlapping odor tunings for ePN and iPN inputs suggest possible counter-balancing interactions that could theoretically implement a high-pass filter (Parnas et al., 2013) in this local region (when ePN input > iPN input). However, in other lateral horn regions, the iPN and ePN odor response tuning mismatched. Understanding how such mismatched feed-forward excitation and inhibition interact and to carry out what computations would need further examination.

### Temporal organization of ePN and iPN responses

In addition to the spatial reorganization of activity, our results indicate that the odor-evoked responses were dynamic and evolved over time at the level of sensory neurons and in both ePNs and iPNs. The initial responses immediately after the stimulus onset were strong but did not have much discriminatory information. Over time, neural activity patterns evoked by different stimuli became more odor-specific. This decorrelation of odor-evoked responses over time was observed in all three neural populations examined. However, the trends observed (which odor pair became distinct when) observed varied even between the dendritic and axonal compartments of the same neural populations, and between flies. This result indicates that a generic computational function can be achieved in an idiosyncratic fashion in flies, and that the information transmitted to the calyx and lateral horns may be qualitatively different.

The decorrelation result is strikingly similar to what has been reported in other model organisms, particularly in zebra fish (Friedrich and Laurent, 2001), with one caveat. We found that decorrelation already happens at the ORN level and gets accelerated downstream.

However, it is in stark contrast with a recent hypothesis put-forth for odor recognition that suggests initial responses carry information odor identity. One possible explanation for the lack of odor-specificity at the stimulus onset could be that the neural activity immediately following stimulus presentation indicates stimulus presence and help with localization. Such localization signals have been reported in many other sensory systems (Bekesy, 2017). We note that the responses immediately following this localization signal may still be extremely important for the fly to recognize the odorant.

Extraction of odor specific information may happen in two different ways. First, the information may be refined in a systematic manner, such that the initial responses recognize odor groups and additional features are extracted to allow precise recognition (Odor present -> fruity -> tropical -> pineapple; analogous to a decision tree). In this case, a snapshot of activity during later time point is sufficient to recognize the stimulus, while the initial responses may be utilized for other sensory computations. The second possibility is that features are extracted in a serial fashion but the later responses need not be the most unique features. This latter scenario is analogous to serial parsing of words (r·e·a·d· vs. r·e·e·l· vs. r·a·i·l· vs. m·e·e·t·). While the initial letters are still important for word recognition, the subsequent letters extracted are necessary but in isolation are not sufficient to allow precise recognition. In this case, an integration of all the features extracted might be necessary for stimulus recognition. Our results indicate that temporal patterning observed in the fly antennal lobe may be more analogous first scenario (i.e. pairwise similarity smoothly reducing over time), but achieved in an idiosyncratic fashion, indicating multiple different solutions may exist to this problem.

It would important to point out that variations across different individuals could arise trivially due to unaccounted differences in experimental conditions between different experiments. However, our results reveal that not all results we observed varied across individual flies. First, as highlighted earlier, gross spatial features matched across individual flies (**Figure 4**). Further, even in the temporal dimension, certain pairs of odorants evoked responses that were highly consistent (**Figure 7**). Such robustness in spatial and temporal features, at least for a subset of odorants, indicate that the variations observed in our dataset cannot be attributed solely to trivial differences in experimental conditions.

Finally, our results indicate that the stimulus-evoked responses do not stop after stimulus termination. At the level of sensory neurons, these are persistence of activity, in some cases excitation and other inhibition, that was observed during the stimulus. However, in the ePN and iPN dendrites and axons, the responses often switched from one ensemble to another. Therefore, stimulus ON and OFF responses were orthogonal to each other, and was observed in all flies. These results are consistent with those reported in other sensory systems, and in particular the locust olfactory system (Nizampatnam et al., 2018).

What is the purpose of these elaborate OFF responses? In cockroaches, such responses were observed directly at the level of sensory neurons and were thought to indicate reduction in stimulus concentrations (Burgstaller and Tichy, 2011). Such dedicated ON and OFF neurons were not found in flies. A single ROI in any region was able to respond during either ON or OFF periods depending on the odor. In a different study, it was reported that these OFF responses may indicate ‘unsensing’ of a stimulus (analogous to a pause after a tone or space after word), and were found to be better predictors of termination of behavioral responses (Saha et al., 2017). Furthermore, our results here indicate that the response patterns observed after stimulus termination were stimulus specific and more diverse than those observed during the stimulus presence period. Further, when odorants are encountered in sequences, the OFF response of the first stimulus was found to contrast enhance the neural activity evoked by the second stimulus. While these results are similar to the findings observed in locusts, causal relationship between OFF responses and their behavioral contributions remains to be determined.

## Acknowledgements

We thank members of the Raman Lab (Washington University in St. Louis) for feedback on the manuscript. We thank Dr. Paul Taghert for kindly providing advice regarding experiments and the lab space for HR to breed flies and perform dissections. We thank Dr. Cody Greer, Dr. Daewoo Kim, and Dr. Donghoon Lee for helping substantially with imaging system troubleshooting and control script development. We thank M.S. Dennis Oakley, Dr. Michael Chien-Cheng Shih and Dr. James A.J. Fitzpatrick at Center for Cellular Imaging (WUCCI) for years of operational support. We thank Dr. Zhen Peng and Dr. Xitong Liang for advising fly breeding and crosses. We thank Dr. Jing Wang (UCSD) for advice on fly preparation. We thank Dr. Tzumin Lee for generously providing *MZ699-Gal4* flies and Bloomington Stock Center for making all other fly strains used in this work available to us. This research was supported by NSF Neuronex grant (Grant# 1707221) and a NSF CAREER Award (Grant #1453022) to B.R.

## Author contributions

HR and BR conceived the study and designed the experiments/analyses. HR performed all the light-sheet experiments and collected the data. LZ motion corrected the raw calcium imaging movies and extracted ROIs in collaboration with HR. HR performed all the subsequent data analysis with LZ’s assistance. HR, LZ and BR wrote the paper taking inputs from all the authors. YBS and TH advised on the selection of fly lines and light-sheet imaging, respectively. BR supervised all aspects of the work.

## Methods

### Fly strains and culture conditions/Fly stocks

Flies were raised on a standard cornmeal diet. Vials were kept at 25° with 12h:12h light-dark cycle. Females 2~6 days after eclosion were used for experiments.

The following fly genotypes were used:

A series of crosses were conducted among

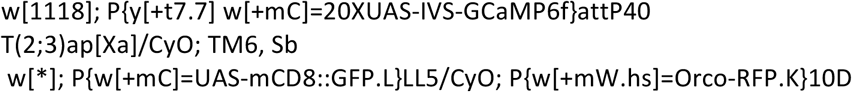

and their hybrid progenies to obtain UAS-GCamp6f; Orco-RFP flies, which were used for crosses with the olfactory neuron tagging GAL4 lines respectively:

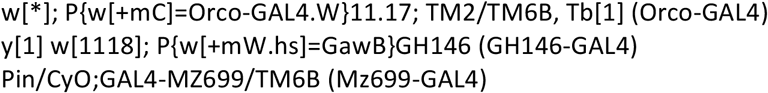

The resulting progenies expressed GCamp6f under the control of neuronal-population-specific drivers (Orco for ORNs, GH146 for ePNs, Mz699 for iPNs) along with RFP expressed in Orco neurons.

### Dissection procedure

The fly was cold-anaesthetized and tethered onto a custom made plexiglass block modified from (Silbering et al., 2012). The antennae were kept underneath the tape film, exposed to the air flow, while the dorsal side of the fly head was immersed in external saline containing (in mM): 103 NaCl, 3 KCl, 5 N-tris(hydroxymethyl) methyl-2-aminoethane-sulfonic acid, 8 trehalose, 10 glucose, 26 NaHCO_3_, 1 NaH_2_PO_4_, 1.5 CaCl_2_, and 4 MgCl_2_ (osmolarity adjusted to 270-275 mOsm) (Badel et al., 2016; Jeanne et al., 2018). Dorsal cuticle was removed. Trachea and stray tissue were cleaned with 5sf forceps (Fine Science Tools). Muscle 16 was cut to stabilize the brain.

### Odor Stimulation

Chemicals were diluted in paraffin oil. In each odor bottle, 20 ml of diluted odor solution (vol/vol) was added. Each batch of odor stimulus was used for no more than 10 days. Stimuli were delivered via a custom-made 16-channel olfactometer. Control signals to the solenoid valves were coupled with microscope control signals. For all experiments carried out in this study, the odor stimulus was 4 s in duration. The onset of an odor stimulus was aligned with the onset of an image stack acquisition. The main air tube was directed at the fly, about 2 cm from the fly. A funnel connected to a vacuum line was placed about 5 cm from the fly block to remove odor residuals.

Stimuli were presented in blocks. The first block comprised of 2~5 trials, during which only spontaneous activities were recorded. To minimize the adaptation and stimulus history related interference that arose due to high neural activities, the odor panel at the lower dilution (10^−4^ v/v) stimuli were pseudorandomized and presented first. Inter-block interval was a minimum of three minutes. Subsequently, a block of odor presentations where each stimulus was delivered at the higher concentration (10^−2^ v/v). The odorants were delivered in the same sequence in both low and high concentration blocks. Then, we again alternated between the low and high concentration blocks at least once more in each fly. Typically, the inter-trial interval within in block was 1 minute. However, for few odorants like 1o30, larger ITI (~1.5 – 2.5 min) were given after stimuli to minimize their interference on the subsequent trial.

### *In vivo* light-sheet imaging

A custom-made light sheet microscope (Greer and Holy, 2019) was used to record imaging data. The microscope has two channels, which we used for recording GCaMP6f and RFP signals simultaneously.

We imaged at ×20 magnification, which provided sufficient resolution for reliable identification of the target neural structures. The typical image size was 1260 × 60pixels, with pixel size being 0.325μm × 0.325μm. The centers of neighboring planes are about 8 μm apart on average. Note each “plane” is in fact a thin volume, as the light sheet kept sweeping through the tissue during the short camera exposure. At most 1 μm (upper bound) may be missed between two optical planes, which is smaller than the neural structures of interest.

Each brain volume was sampled at 4 Hz. For ePN and iPN recordings, a volume of ~190 μm thickness were scanned through along the axis of piezo movement (z-axis) to cover both the antennal lobe and calyx/lateral horn as these regions reside in different optical planes. The data from iPN dendrites were discarded from subsequent analysis due to the extremely low GCamp6f signal in the region. Calcium signals from a volume of ~80 μm thickness was recorded for monitoring responses at the ORN axonal terminals in the antennal lobe.

488 nm and 561 nm lasers were used to excite both the GCamp6f and RFP, respectively. The timings of the two lasers were synchronized to ensure the RFP images were acquired at the same time instances as GCamp signals. The lasers were only turned on during the camera exposure to reduce photobleaching.

During imaging experiments, external saline oxygenated with 95%O_2_/5%CO_2_ (Airgas), was perfused at 2mL/min. Only flies that showed some change in calcium signals during a paraffin oil puff or 10^−4^ odor test pulse were chosen for formal recording. Data acquisition began at least 5 min after the end of the test pulse.

### Motion correction

We pre-processed the imaging data to correct for motion artifacts during acquisition. As the functional imaging movies generally contain flashing activities of neural response, it’s difficult to obtain static reference template. We found that most of the motion artifacts, if present, in our datasets were due to translational displacements. To remove these artefacts, we used two different strategies to account for motion artifacts in the antennal lobe and in the lateral horn. In antennal lobe, we first corrected the motion using simultaneously acquired anatomical imaging data (RFP labeled ORN axons). Using the anatomical dataset, we found the translation correction matrix that maximized the correlation value between the target frame and template frame. Then we used this translation matrix to function recordings in the antennal lobe and obtained motion corrected imaging data. In the lateral horn, we learned the translation matrix by focusing on the less responsive regions (neural tracts). The obtained translation matrix was used to correct for the overall motion artifacts in the whole image.

### Identifying response regions of calcium imaging data

We identified regions of interest (ROIs) by applying a constrained nonnegative matrix factorization (Pnevmatikakis et al., 2016). The spatiotemporal calcium activity can be expressed as a product of a spatial basis matrix A and a temporal matrix C.

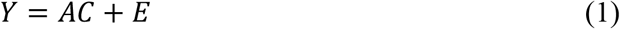

*Y* represents spatiotemporal calcium responses, where each column represents vectorized calcium image in a time frame and each row represents a pixel value across time frames, and *E* indicates the observation noise. The factorization procedure is similar to regular nonnegative matrix factorization, requiring spatial matrix *A* and temporal matrix *C* being nonnegative. Moreover, the spatial component matrix is endowed with additional sparsity constraint to extract more compact and regularized spatial response regions as ROI masks. The problem can be succinctly summarized as the following optimization problem:

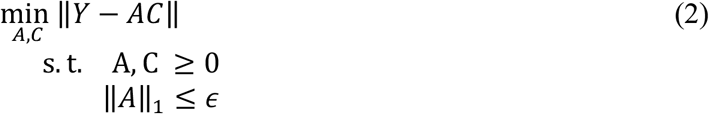

We optimized the spatial component and temporal component by alternating such that a new estimate of *A* is obtained by use of the last estimate of *C* and vice versa. As both subproblems are convex, there exists a variety of methods to solve it. We solved the spatial subproblem by a nonnegative least-angle regression (LARS) algorithm and temporal subproblem by nonnegative least squares. We used different degrees of spatial constraints (*ϵ*) to account for various responses statistics in antennal lobe and lateral horn. Similar to (Pnevmatikakis et al., 2016), at the end of each iteration, we merged overlapping components with high temporal correlation and removed components with low signal-to-noise ratio.

### Initialization of CNMF using local correlation map

Even though the individual sub-problems are convex, the overall optimization problem listed above is non-convex. The quality of solution is highly sensitive to the initialization. Exploration of initialization methods is time consuming and computationally expensive. Additionally, often requires a preset number of spatial components need to be identified during initialization (i.e. number of columns of matrix A). In this study, we used a local correlation map based approach to initialize the response regions (i.e. matrix A). The correlation value in each pixel is obtained by computing correlation coefficients between the temporal trace of that pixel and the mean temporal trace of surrounding four pixels (i.e. one above, one left, one right and one down). After we obtain the local correlation map, we apply a median filter and morphological closing to obtain the initial response regions (columns of matrix A). Compared to other initialization methods, this approach was computationally more efficient and the number of spatial components required for factorization was automatically determined based on the imaging dataset.

### Correction of exponential signal drifts within each trial and calculation of *ΔF/F*

First, the camera bias, a constant value, was subtracted from all signals acquired. A robust estimation of the baseline F at each time instance is essential to the reliable calculation of ΔF/F. However, three common phenomena made this task challenging: 1. Intra-trial baseline drift, an approximately exponential decay within each trial (intra-trial drift), possibly due to photobleaching. 2. inter-trial baseline drift, the baseline may drift between trials, possibly due to some slower cellular processes. 3. Noisy spontaneous signals, instead of a “real baseline”, the observable signals are the results of random fluctuations or spontaneous activities being superimposed on the underlying baseline.

To tackle these problems, we devised an approach to model the spontaneous fluorescence signals based on two basic assumptions: 1. The “true baseline” underlying the observed signals is a constant value for a given trial. Meanwhile, given the inter-trial baseline drift, the “true baseline” is trial dependent. 2. The observed signals are a result of superimposing an exponential decay on top of the “true baseline”, and the rate of this exponential decay for a given ROI is fixed.

Hence, the observed spontaneous signal *F*_*t*_*′* at time instance t in a trial can be described by formula

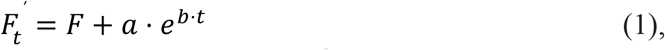

where F is the “true baseline” of the given ROI in that trial, and *a* · *e*^*b · t*^ is the exponential term describing the intra-trial signal decay. To remove the contribution of uncorrelated noise observed in different trials, the exponential term was modeled as the “mean” exponential decay of all the trials for a given ROI.

To obtain the data for exponential term estimation, for each ROI, we pooled the pre-stimulus signals (first 2.5 s excluded) and the very last 1 s from each trial, which resulted in a *m*×*n* matrix, with ‘*m*’ indicating the total number of trials and ‘*n*’ indicating the total number of sampled time points. We computed the standard deviation of each column. Values out of the ±1.5 std range in the column are discarded as outliers. Then we parameterized the formula

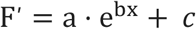

by fitting it to the remaining data points within the pool while minimizing the mean squared error (MSE). Note, at this step, our goal was to obtain the mean intra-trial exponential decay term. Constant c describes some sort of the ROI’s “mean baseline” across trials. Now for a given trial we have the “true baseline”

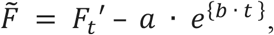

where the exponential term is already known. Next, to obtain a trial’s F, we simply parameterize 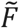 by minimizing the MSE between formula (1) and that trial’s spontaneous signals.

The baseline correction approach resulted in a small fraction of ROIs having near zero or even negative baselines values. Since this could result in unrealistically large ΔF/F, we dealt with this issue in the following fashion.

To minimize the amount of baseline correction, these ROIs also had to meet a set of criteria to ensure it’s ΔF and ΔF/F are indeed outliers of the population and the baseline value must be under an empirically determined threshold. For such an ROI, we substituted its baseline with the mean baseline across all other ROIs on the same plane. Note that in cases where the corrected baseline was smaller than the original baseline *F*_*i*_, the original baseline was retained.

### ROI cleaning

The ROI masks were projected back to the raw image movies. ROIs that does not belong to the target structures were removed after visual inspection.

Given some ROIs in the antennal lobe can span more than one planes and possible errors made by the detection algorithm, we sought to remove the duplicates. Candidate duplicate ROIs were identified by running a hierarchical clustering analysis on following response features:

- cosine distance between high-dimensional vectors of calcium signals recorded during this whole trial [34.5 s total: −9.5 s before odor onset to 25 s after odor onset; ~ 138 dimensional vectors]
- cosine distance between calcium signals recorded during a 15 s post-stimulus period [10 s after stimulus onset to +25 s after odor onset; ~ 60 dimensional vectors]
- Euclidean distance between a 15 s post-stimulus time periods across different trials [10 s after stimulus onsest to +25 s after odor onset; ~60 dimensional vectors].

The resulting candidate set was the intersection of the candidate sets generated by the independent hierarchical clustering.

Finally, the candidates were mapped back to the anatomical space, and re-examined through visual inspection. A candidate ROI was labeled as duplicate, only if it were clustered together with another ROI and was anatomically juxtaposed to it.

### Quantification and Data Analysis

#### Quantification of ROI functional distance

An ROI’s response to a stimulus was represented by its mean ΔF/F observed during the odor presentation window. Therefore, for a given ROI, its tuning was represented by a 12-dimensional vector, since the odor panel used in the study included six odorants each delivered at two different intensities [i.e. 12 stimuli]. The functional distance between an arbitrary pair of ROIs (ROI A, ROI B) was defined as the cosine similarity, defined as:

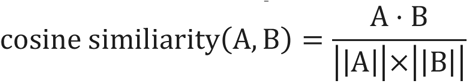

between their 12-D tuning vectors. The spatial distance between two ROI’s was calculated as the Euclidean distance between their centroids in the physical space.

The functional and spatial distances between pairs of ROIs were calculated and pooled across individual flies. The relationship between the two distances was determined using a linear regression. The degree of “linearity” between these two parameters was quantified using the R-squared value of the best-fit linear model, e.g. the amount of variance that can be explained by the model.

### Functional embedding and the projection onto anatomical space

To visualize the relationship between the ROI “tuning” and the spatial organization, one intuitive approach is to represent an ROI as a point using its centroid coordinates in the 3D anatomical space, and assign similar colors to these points that have similar stimulus preference or tuning. Namely, for an arbitrary pair of ROIs, if their functional distance is small (i.e. similar tuning), colors that represent them should be close to each other in the RGB color space as well. Given that the RGB color space is essentially a 3D space, we used multidimensional scaling (MDS) to translate the pairwise functional distance into Euclidean distance in the 3D RGB color space. The pairwise functional distances of all ROI pairs were precomputed as a “dissimilarity” matrix and fed into the parametric MDS algorithm. The resulting 3D coordinates of the ROIs were normalized to unit scale by the following procedure:

Let *X* be the set of all x-axis values of the MDS output. Let *X*_*ceil*_ be the 95% quantile value of *X* and *X*_*floor*_ be the 5% quantile value.

We have the normalized value 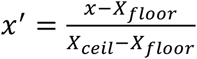. If *x*′ is out of the range [0,1], it was clipped to either 0 or 1, whichever was closer. This procedure was repeated on values corresponding to the other two MDS axes.

Note the max and min values were defined as the values at the 95% and 5% quantiles, respectively, for robustness. Then the normalized coordinate values were used as RGB values. In addition, several landmark tuning vectors were artificially constructed and added to the dataset to facilitate interpretation. Sharing the same 12-dimensional format as the real ROI tuning vectors, these landmark tuning vectors had 1s indicating excitation, 0s indicating no response, and −1s indicating inhibition instead.

To avoid numerical stability issues, ROIs that were barely activated by any of the odorants in the panel were not considered for this analysis. Less than 1% of the ROIs were neglected due to this criterion.

### Regression Analysis of ePN input and output relation

We regarded the AL spatiotemporal response as the input to the regression model, which was a t×n matrix *X*, where n is the number of AL ROIs and t is the total number of time points (note that responses between 0 to +12s in different trials were concatenated to form a super long column vector). The CL and LH responses from the recording were regarded as the target matrix. Thus, the target matrix *Y* was a t×m matrix, where m is the total number of CL and LH ROIs. We have the generic form of linear regression:

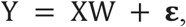

where W is the n ×m weight matrix that transforms AL response into CL/LH response, while minimizing the error ε. Since direct least-squares regression to determine W was not feasible, we used a multi-task lasso regression (MTLR). Optimal W was obtained by minimizing a slightly modified objective function:

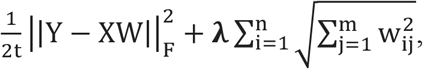

where 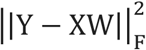 is the Frobenius norm of the residual matrix with 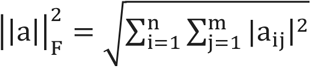 e.g. the square root of the residual sum of squares of each element. Note the regularization term is essentially a *l*_1_-norm of *l*_2_-norms, scaled by the hyper-parameter λ.

To determine the optimal hyper-parameter, we performed a grid search, adopting a K-fold cross-validation scheme that leaves one stimulus group (both concentrations of the same odorant) out each time.

### Analysis of temporal coding

To quantify the pattern similarity between a stimulus pair as a function of time (**Fig. 6**), we first aligned trials with respect to stimulus onset. Then we computed the cosine similarity between the two population response vectors at the same reference time point (i.e.cosine (a_t_,b_t_) where a_t_ and b_t_ are the at time *t* following introduction of stimulus a or b, respectively). By computing the similarity at different points in time after odor onset, we characterized how similarity between pairs of odorants evolve as a function of time.

Since the odor panel comprised of six odorants each delivered at two concentrations, we calculated similarity between 66 unique stimulus pairs in total.

For the visualization of cosine similarity distributions, kernel density estimation was performed using a Gaussian kernel with the bandwidth determined by the Scott rule (Scott, 17 August 1992).

### Analysis of ON-OFF response

The neural activities during the 4 s stimulus presentation period was defined as the “ON response,” whereas activities during a 4 s time window after the stimulus termination were taken as the “OFF response”. The one second period immediately following the termination of the odorant was excluded as it included both ON and OFF responses.

We used a MDS dimensionality approach to visualize the ON or OFF response vectors (Supplementary Fig. 11). The MDS analysis was done independently for data collected from each individual fly.

To quantify the diversity of the ON and OFF response patterns, principal component analysis was performed on the same data. The number of principal components (PCs) needed to account for at least 90% of the variance in activity patterns was used to measure the pattern diversity, as more diverse patterns would require more PCs to capture the majority of the data variance, and vice versa (**Fig. 8D**).

To compute the mean angle between the ON and OFF activity patterns, for each stimulus, mean activity pattern vectors were computed for the ON and OFF time windows. For a given stimulus pair, the angles between the ON and OFF activities were calculated and averaged across individual flies (**Fig. 8C**).

## Supplementary Figure Captions

**Supplementary Figure 1:**
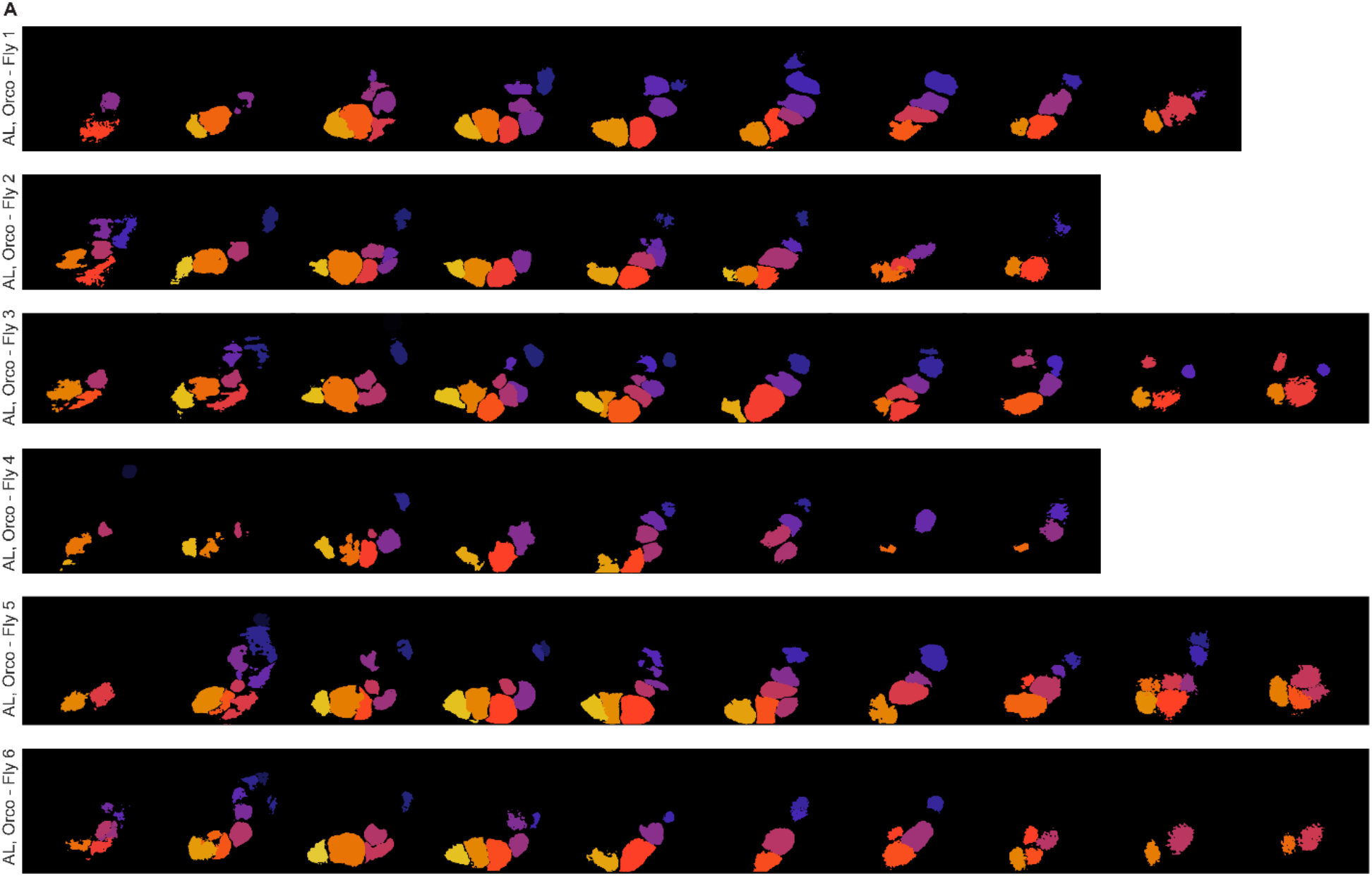
ROI masks extracted to segment axonal responses in flies expressing calcium indicators in ORN axons. ROI masks extracted for each plane and in each Orco labeled fly are shown. Each row shows ROIs across different planes for an individual fly. Left most panel shows ROI masks in dorsal regions and right most panel shows ROIs in more ventral regions. In each plane, different ROIs are labelled using different colors. In total, ROI masks for all six flies used in the study are shown in different rows.

**Supplementary Figure 2:**
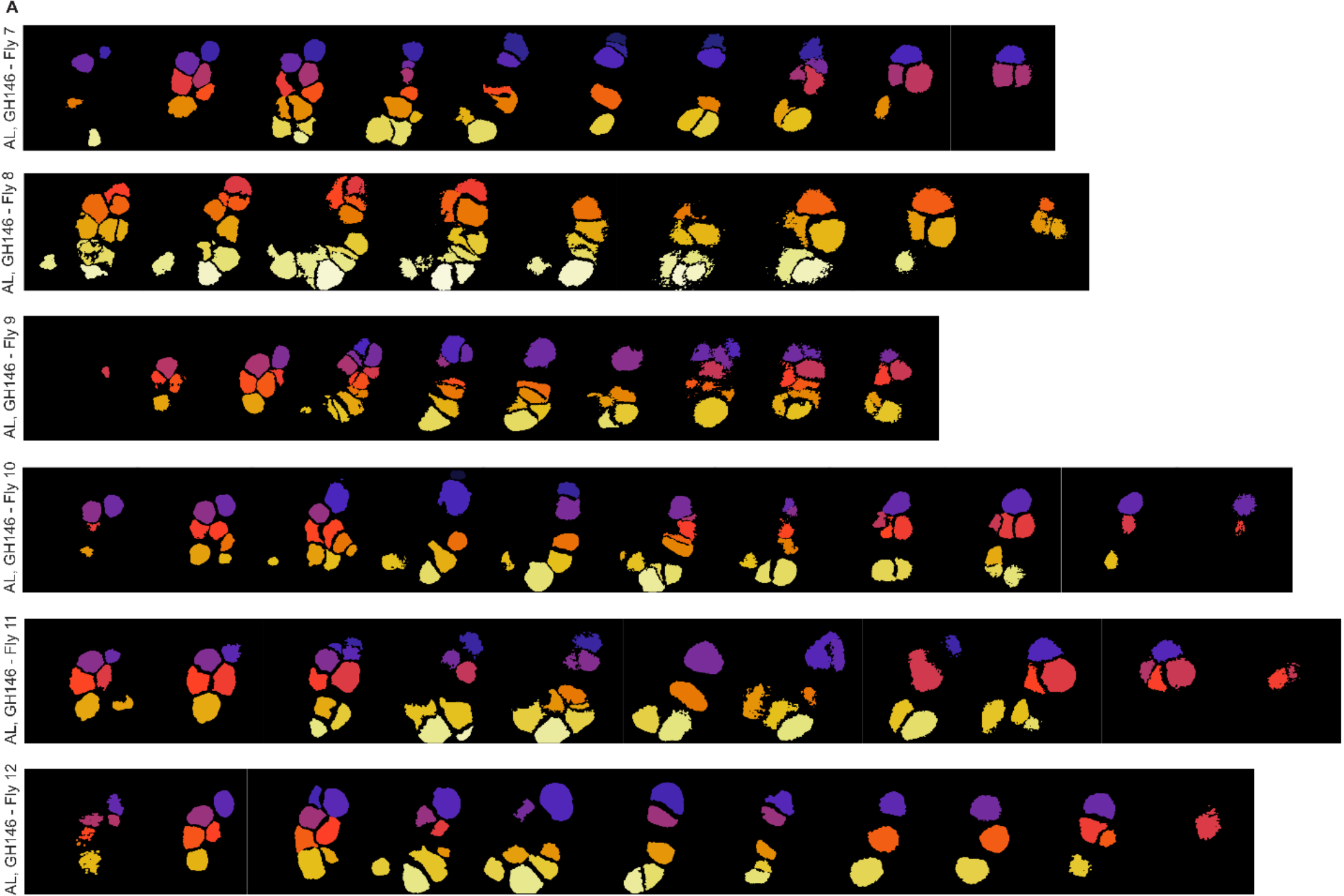
ROI masks extracted that segment ePN dendritic activity in the antennal lobe. Similar to **Supplementary Figure** 1, but ROI masks for GH146 flies with ePNs labeled with GCamp6f. ROI masks extracted are shown for each plane and in each fly antennal lobe i.e. to segment dendritic responses.

**Supplementary Figure 3:**
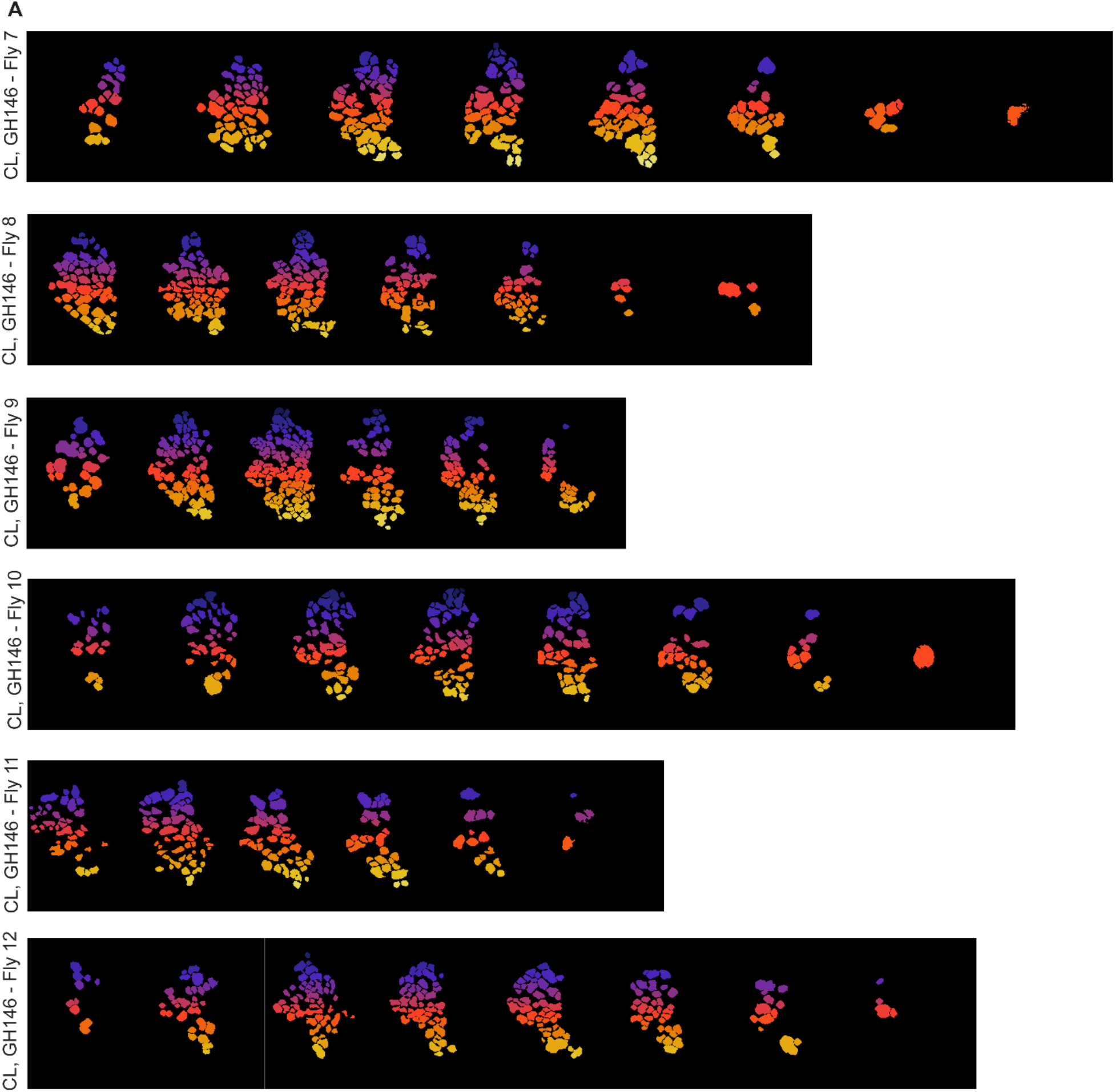
ROI masks extracted to segment ePN axonal responses in the mushroom body calyx. Similar to **Supplementary Figure** 1, but ROI masks to segment ePN axonal responses that are transmitted to the mushroom body calyx are shown for each plane and in each fly.

**Supplementary Figure 4:**
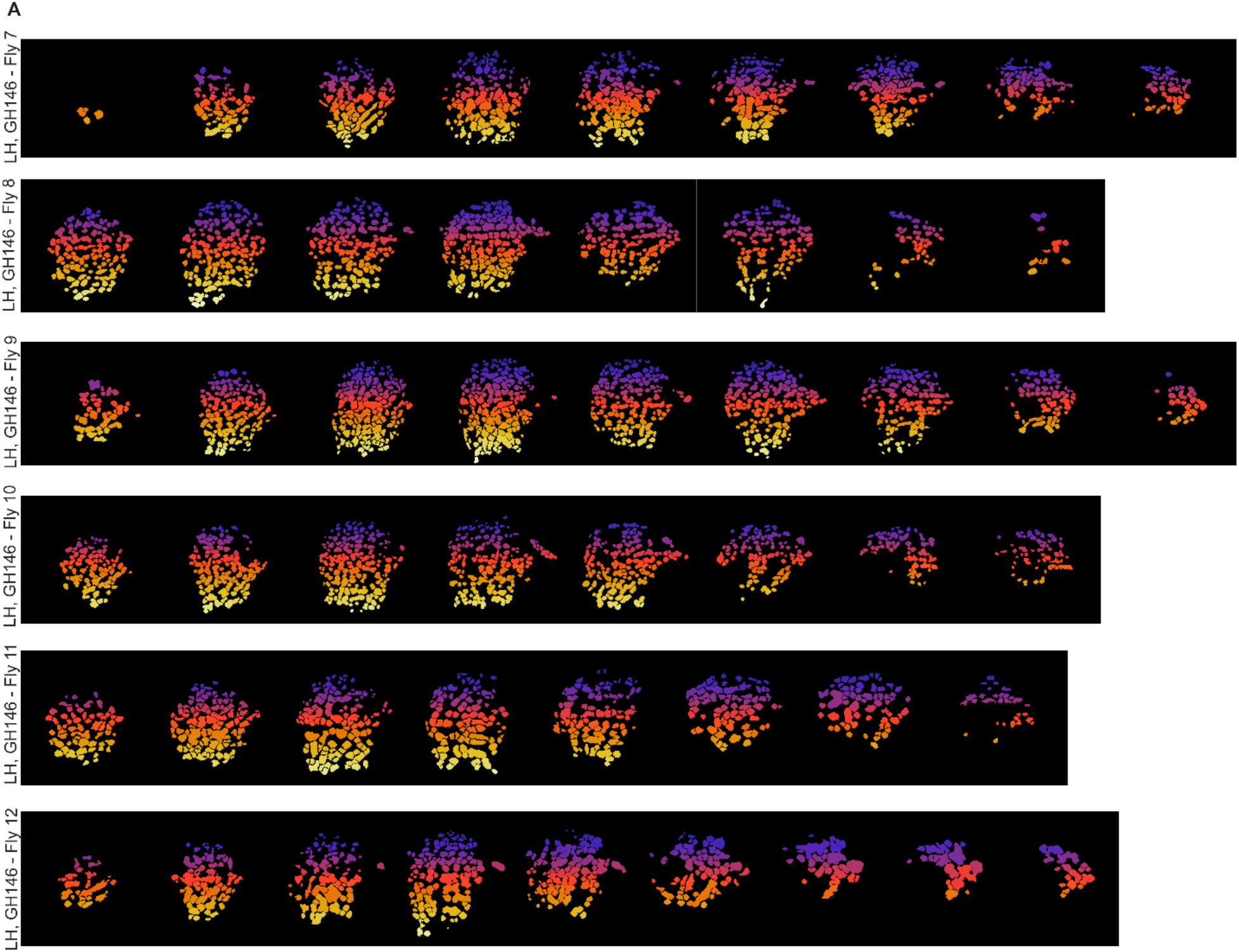
ROI masks extracted to segment ePN axonal responses in the lateral horn. Similar to **Supplementary Figure** 1, but ROI masks extracted to segment GH146 ePNs axonal responses in the lateral horn are shown for each plane and in each fly.

**Supplementary Figure 5:**
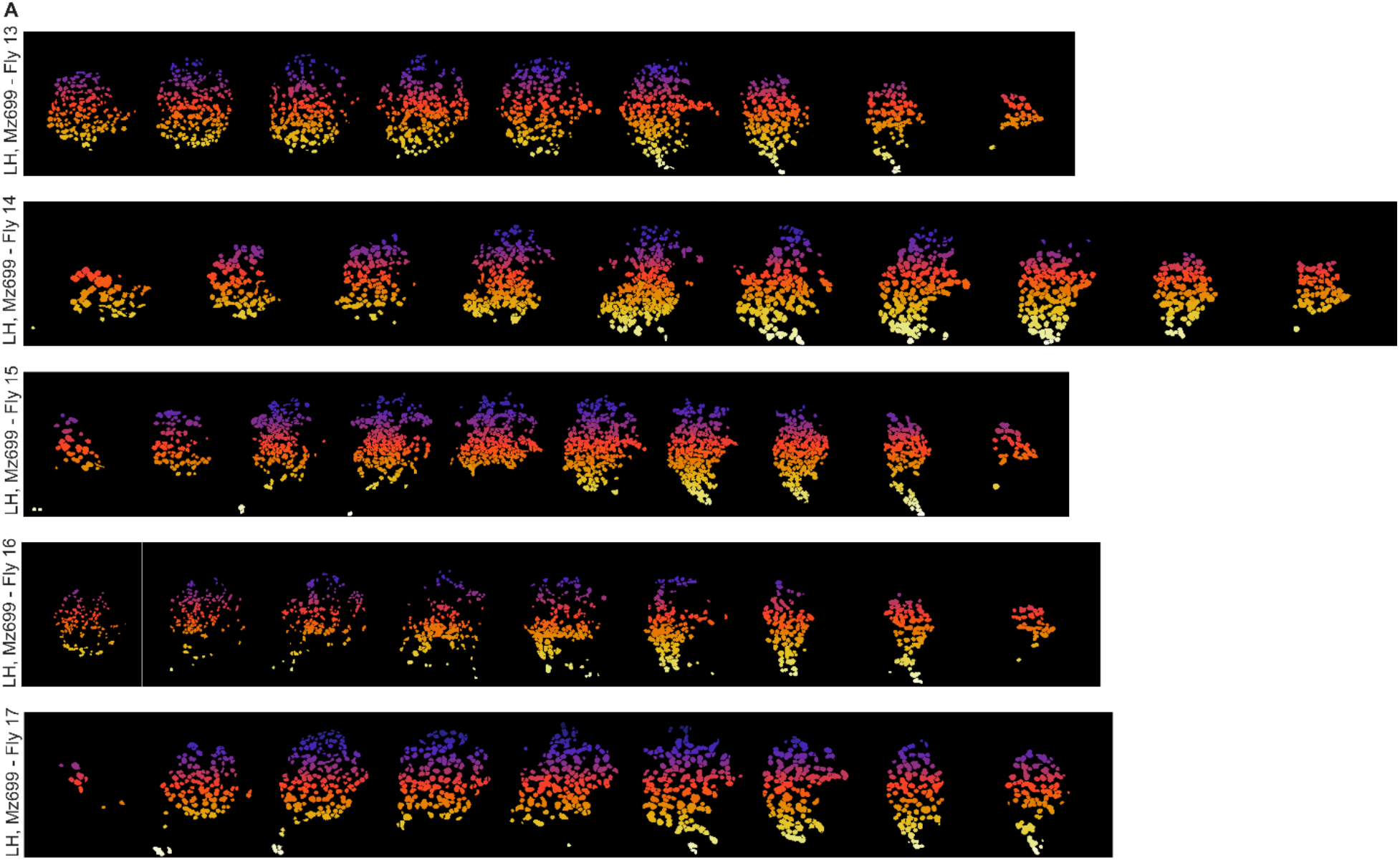
ROI masks extracted to segment iPN axonal responses in the lateral horn. Similar to **Supplementary Figure**1, but ROI masks extracted to segment Mz699 iPNs axonsal responses in the lateral horn extracted are shown for each plane and in each fly.

**Supplementary Figure 6:**
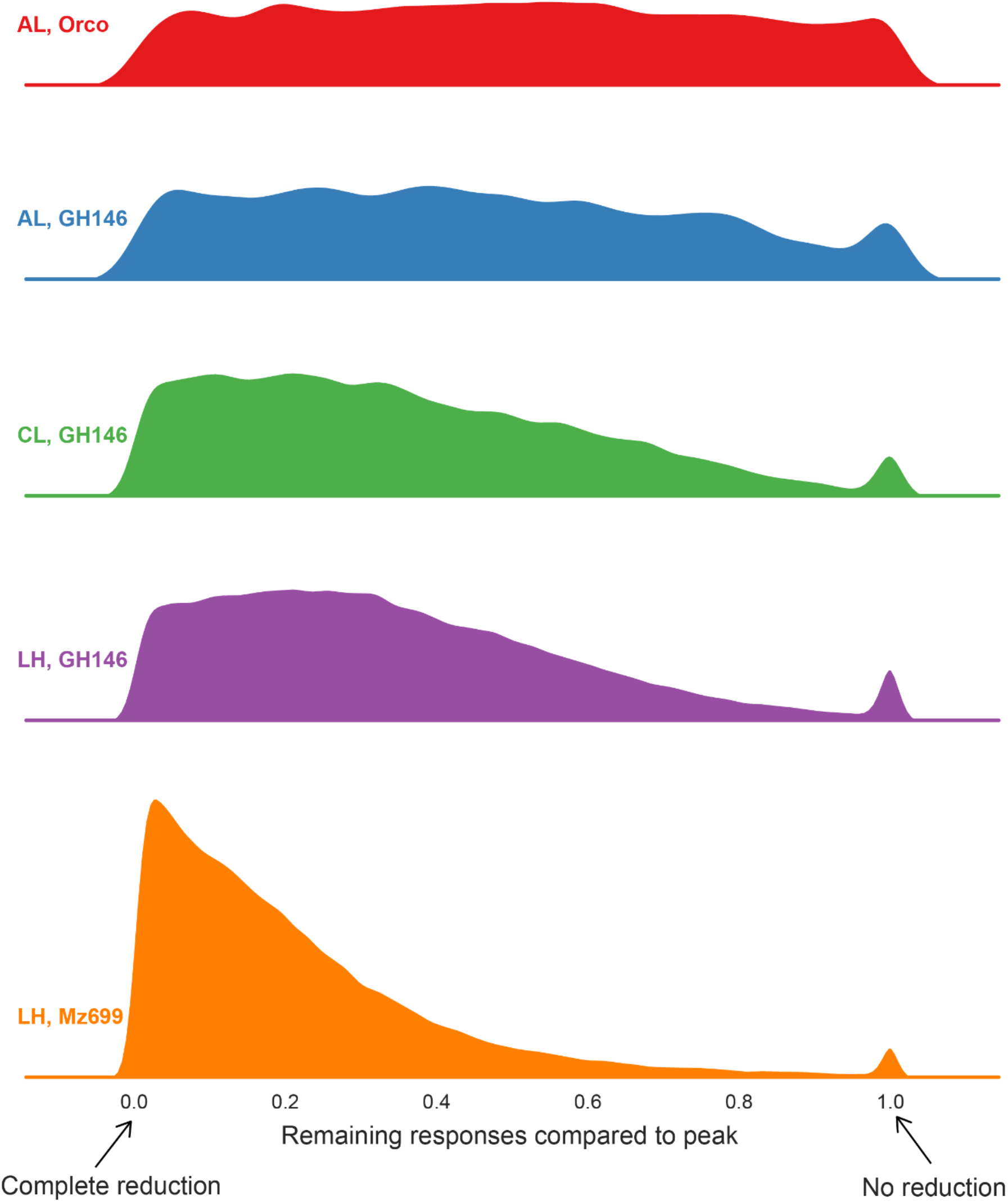
Odor-evoked temporal response dynamics. The response distribution showing the activity levels of ROIs relative to their peak responses at the end of the odor pulse (i.e. prior to termination; 3.75 s after odor onset). To account for excitatory and inhibitory responses, the absolute values of the signals were used for this analysis. The peak response is defined as the maximum absolute value during the odor presentation window. Each row is one fly line/region. The x-axis indicates the fraction of ROIs showing a particular level of activity and y-axis indicates the density.

**Supplementary Figure 7:**
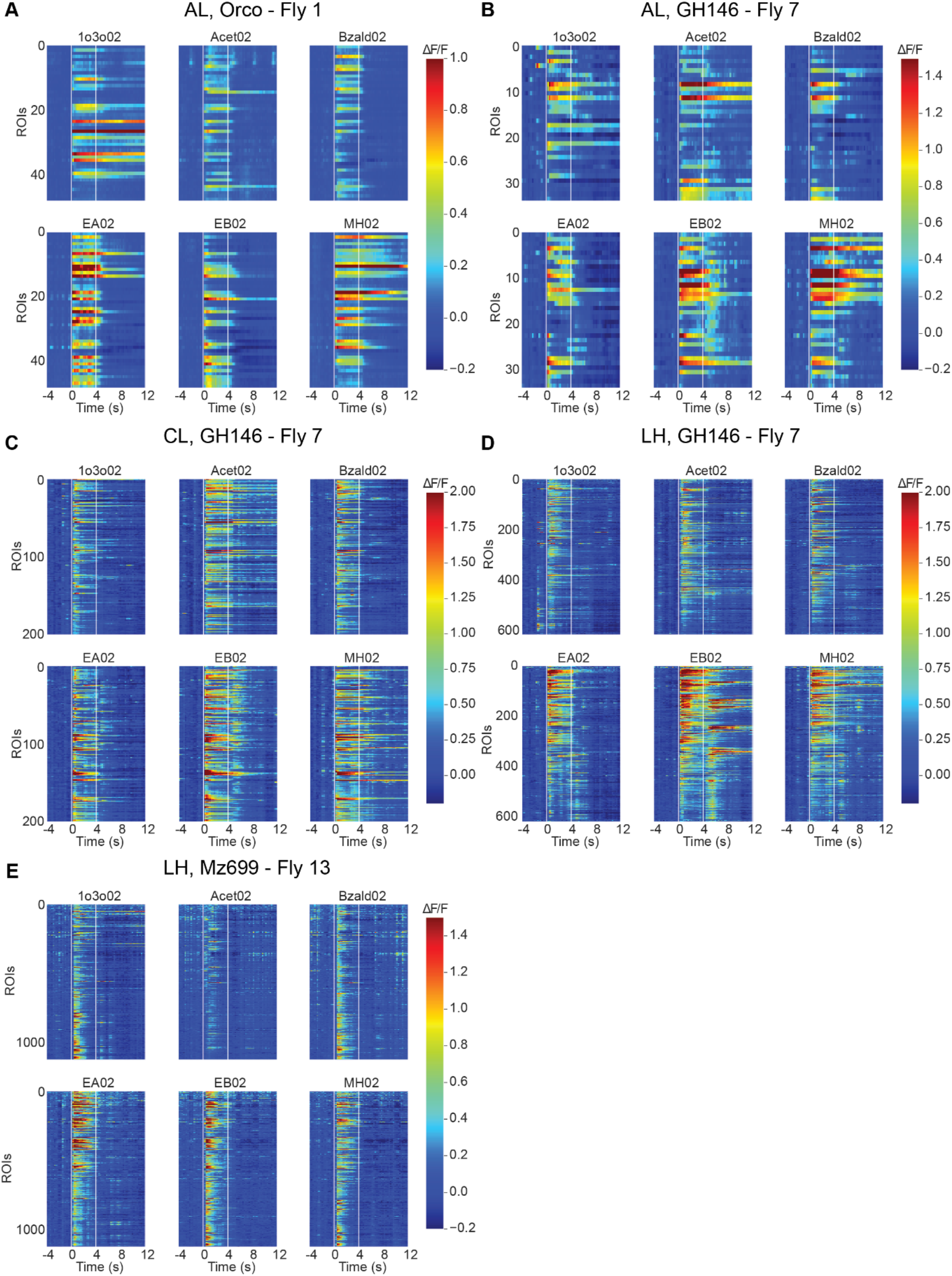
Temporal responses in higher concentrations. **(A** to **E)** Similar to **Figure 2 (B** to **F)** Representative responses to the same six stimuli but delivered at a higher concentration are shown.

**Supplementary Figure 8:**
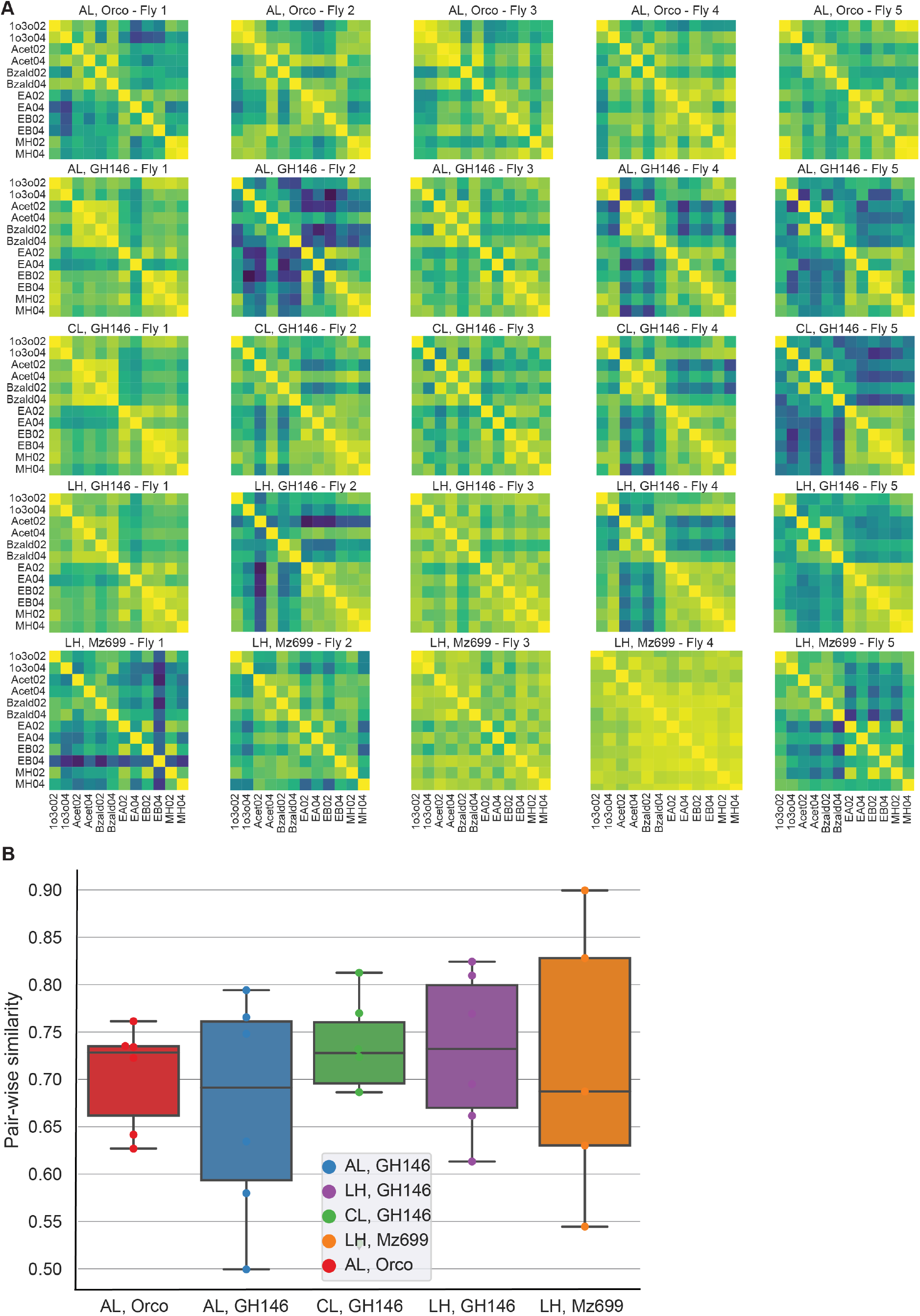
Pair-wise odor response similarities.

**(A)** Representative heatmaps showing similarity between odor-evoked responsese evoked by all stimulus pairs. Each row is a fly line/region and different columns correspond to different individual flies. In each heatmap, each grid is the cosine similarity between a pair of stimuli indicated by the corresponding labels. To calculate the similarity between a stimulus pair, every ROI’s response was represented by its mean response during odor presentation, and the responses across all ROIs were regarded as a high-dimensional vector. The cosine distance between these two response vectors evoked by the two odorants were computed and plotted as a heatmap. Warmer color means higher similarity.

**(B)** Mean pairwise similarities between odorants was computed for each fly line/region and summarized as a box plot. The y-axis indicates the mean cosine similarity between a pair of odorants.

**Supplementary Figure 9:**
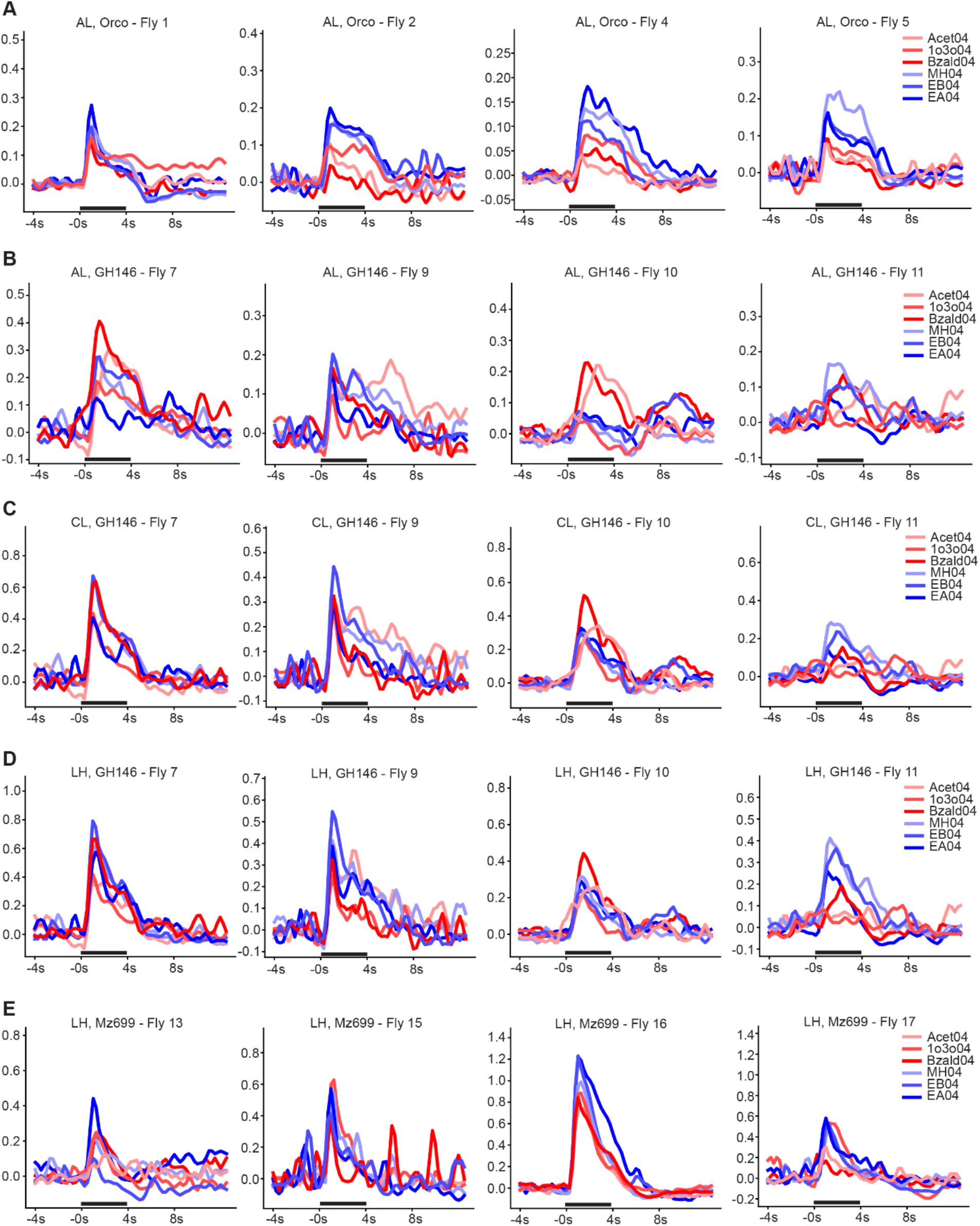
PSTHs characterizing overall responses evoked by the odor panel at lower concentration. **(A)** Mean firing rates across all ORN axon ROIs are shown for four representative flies. In each panel, responses to 6 different stimuli delivered at their lower concentrations are shown. Red color are used to indicate PSTHs evoked by putative repulsive odorants and blue colors label PSTHs evoked by attractive ones. The 4-s odor stimulation period is shown as a black bar along the x axis. **(B)** Similar as panel **(A)**, but mean firing rates across all ePN dendritic ROIs in the antennal lobe are shown. **(C)** Similar as panel **(A)**, but mean firing rates across all ePN axonal ROIs in the calyx are shown. **(D)** Similar as panel **(A)**, but mean firing rates across all ePN axonal ROIs in the lateral horn are shown. **(E)** Similar as panel **(A)**, but mean firing rates across all iPN axonal ROIs in the lateral horn are shown.

**Supplementary Figure 10:**
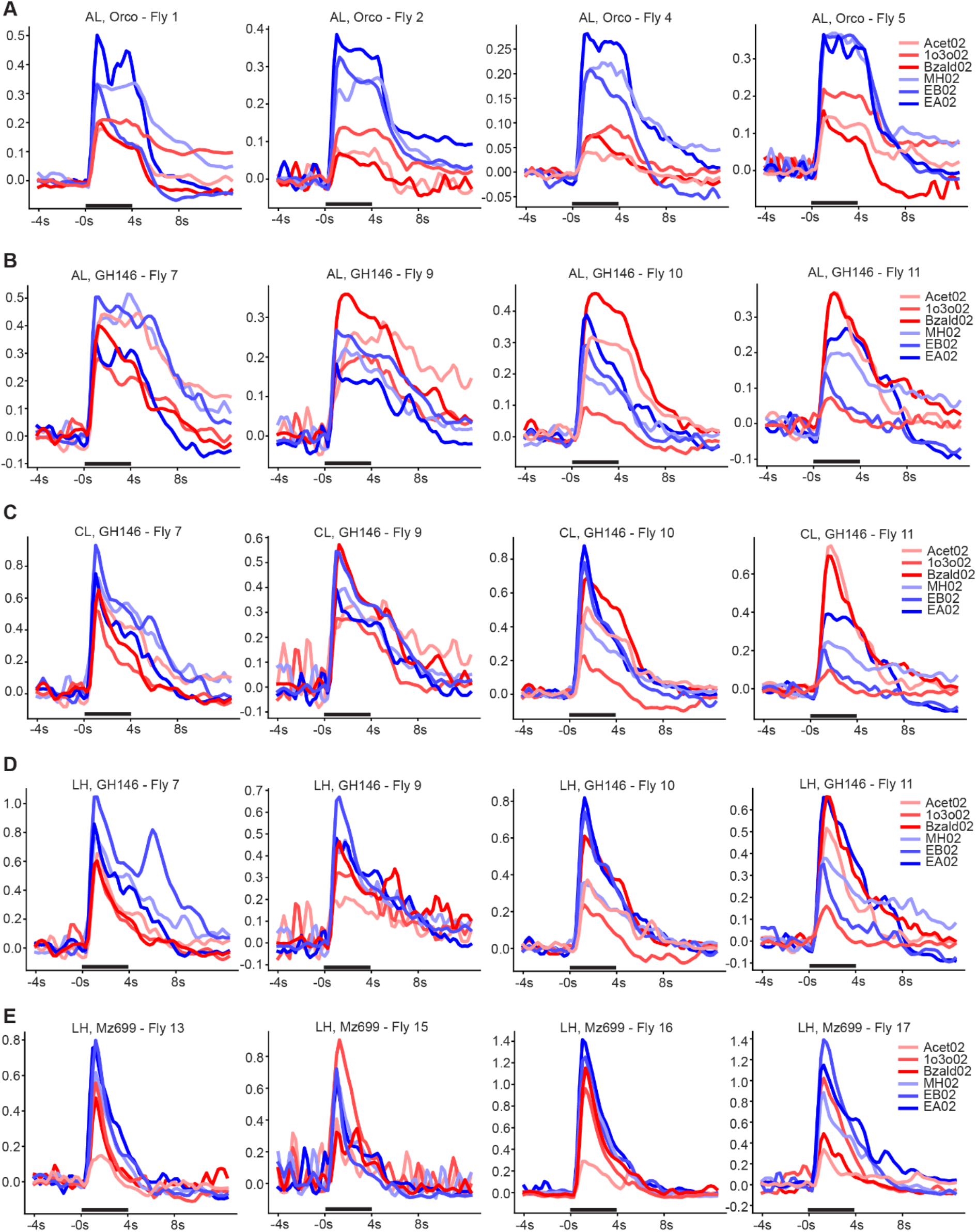
PSTHs characterizing overall responses evoked by the odor panel at lower concentration. Similar as **Supplementary Figure 8**, but mean firing rates for the same panel of odorants delivered at a higher concentration are shown.

**Supplementary Figure 11:**
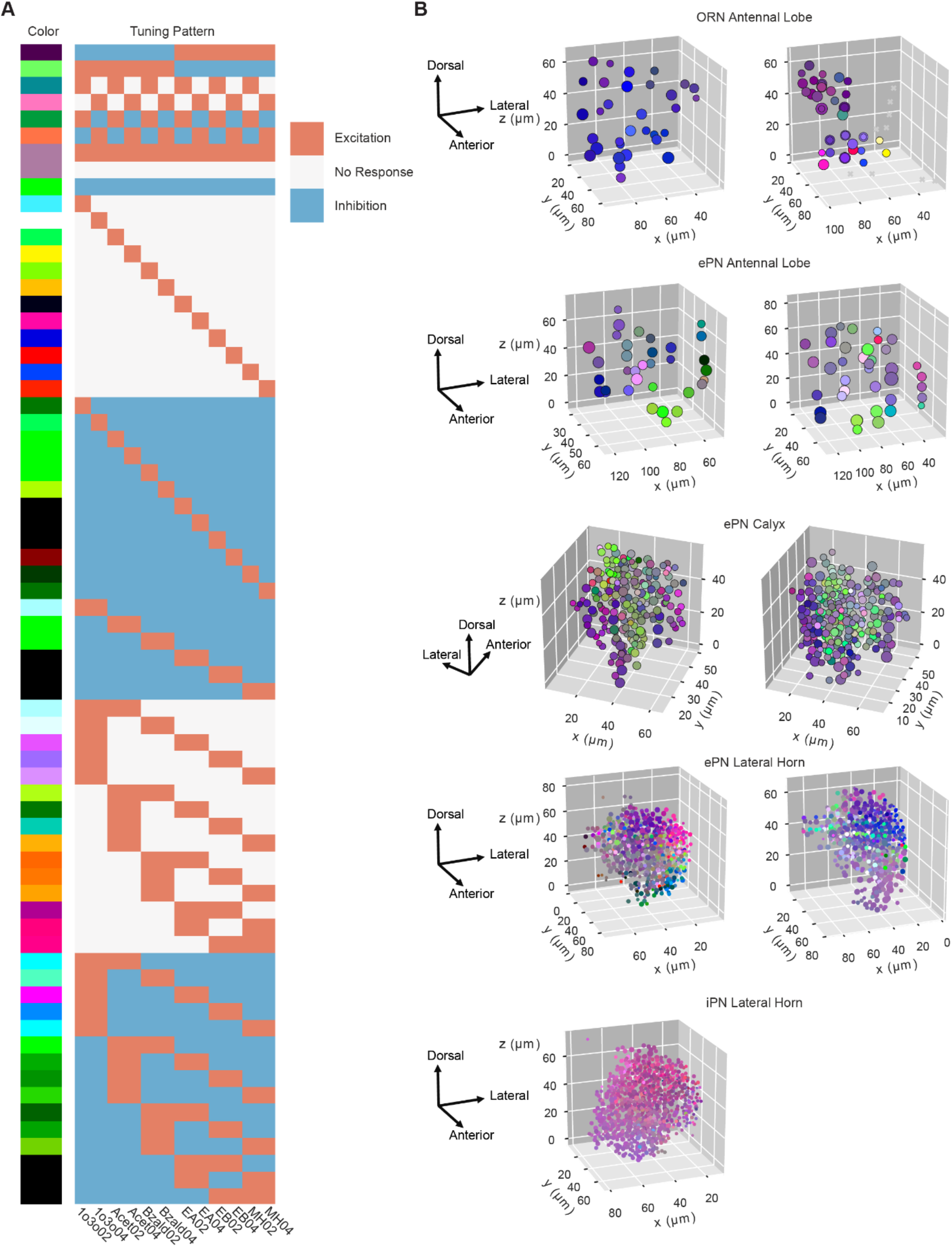
A more elaborate set of reference vectors/templates.

**(A)** The mapping between tuning vectors onto the 3D color space is illustrated with a more elaborate set of reference vectors/templates. Each row corresponds to a mapping between a reference template and the color assgined.

**(B)** Similar to **Figure 4(B)**, ROIs are shown in actual spatial locations across different regions but for two additional flies in each line/region.

**Supplementary Figure 12:**
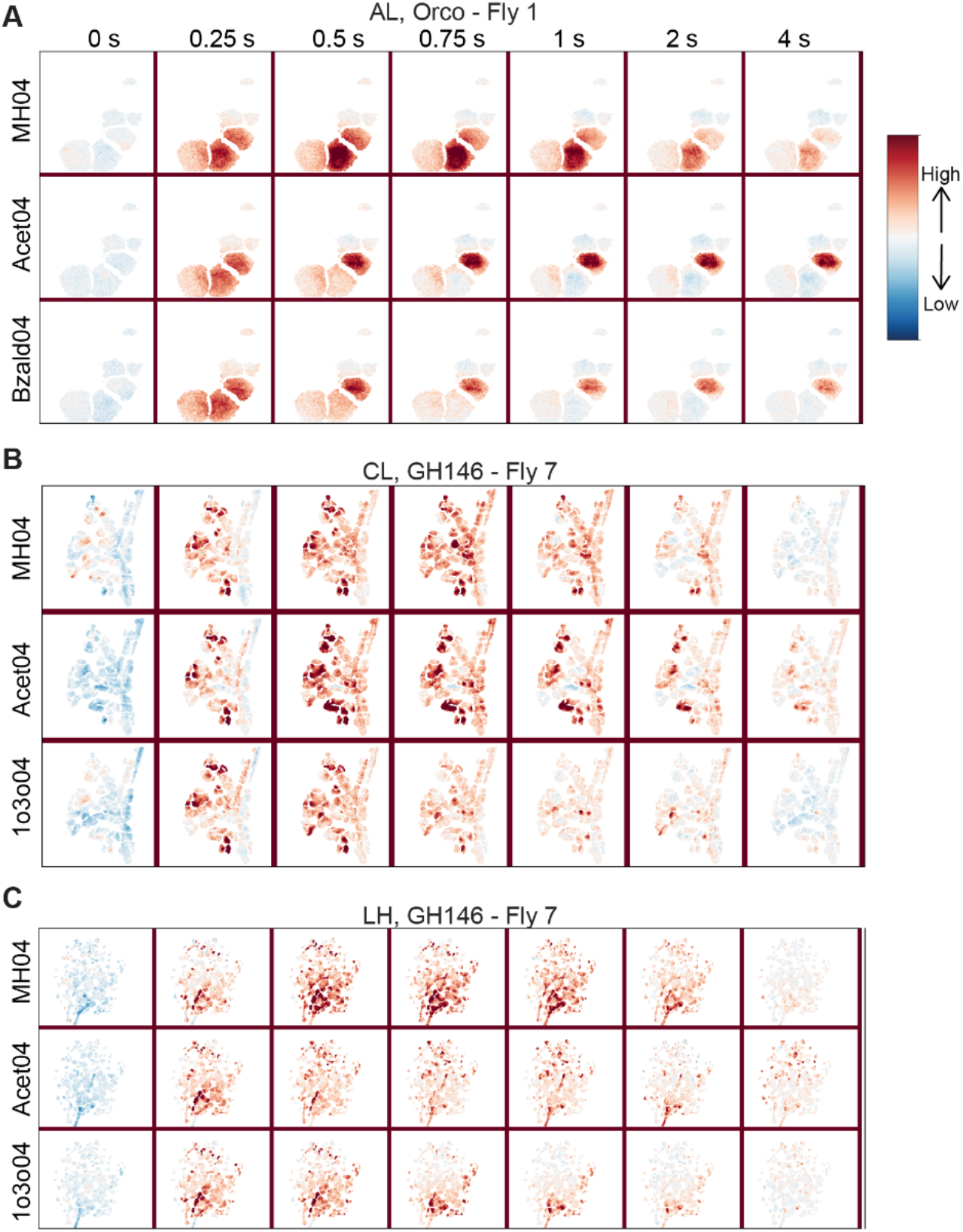
Time evolution of spatial responses in ORNs and ePN axons. **(A** to **C)** Similar plot as shown in **Figure 6(A)**, but showing spatial distribution of activity at the level of ORN axons in the antennal lobe, and ePN axons entering the calyx and the lateral horns. Time since odor onset is shown at the top of each panel: 0 s indicates start of odor stimulus and stimulus lasts for 4 seconds. Response to three representative odorants are shown in the three rows.

**Supplementary Figure 13:**
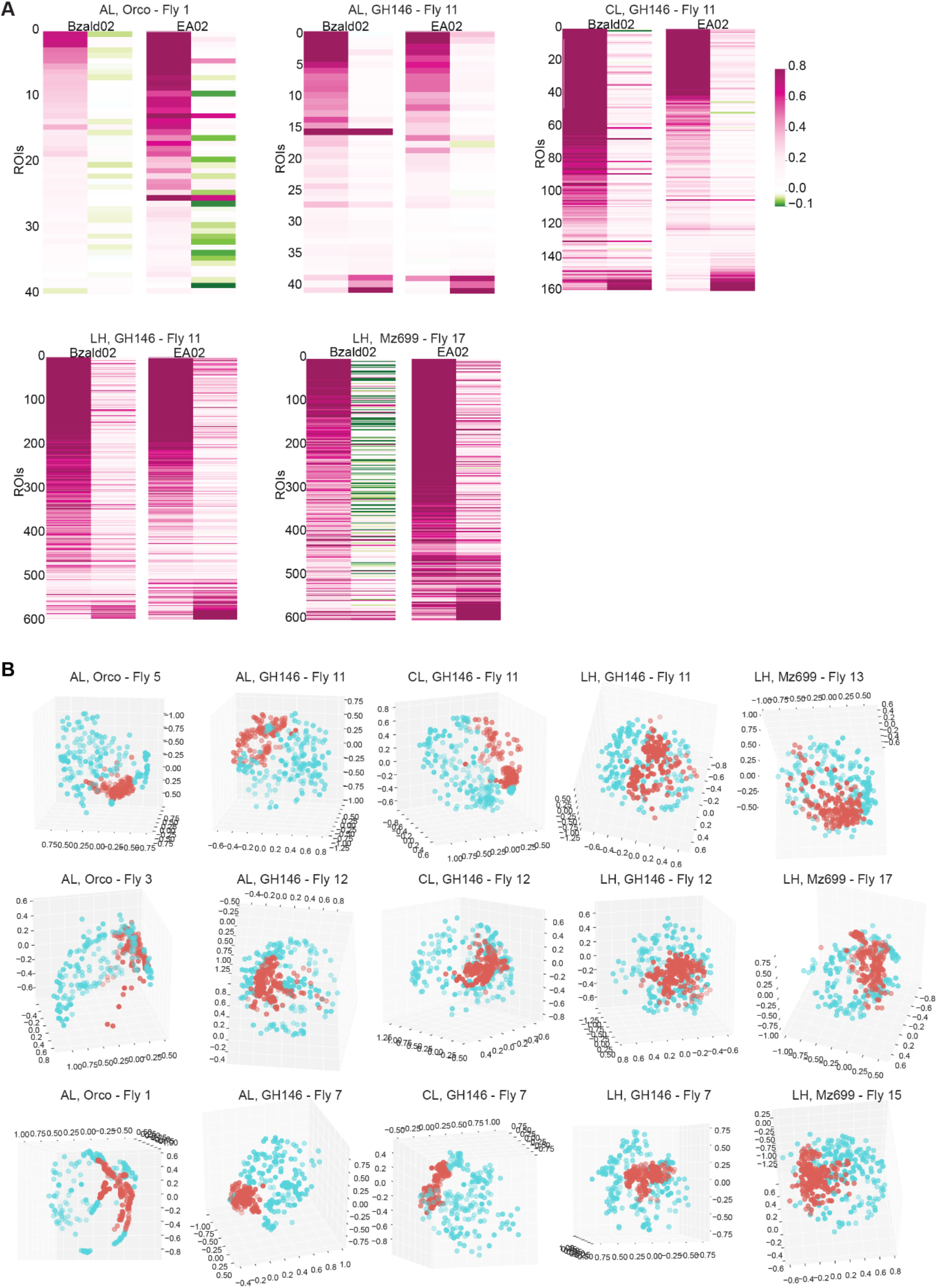
Odor evoked ON vs. OFF responses. **(A)** The average responses across all ROIs during the ON and OFF response periods are stacked next to each other and shown as a color bar. In each panel, left column indicates ON responses (peak activity during 4s ON window) and right column shows the OFF responses (peak activity during 4s window after termination of the stimulus). Each row represents one ROI. **(B)** ON vs OFF response pattern comparison following visualization using a multi-dimensional scaling (MDS) approach. The ensemble responses at each time point during the ON and OFF period were regarded as high-dimensional vectors, and were plotted in a 3D plot after MDS dimensionality reduction. Response vectors evoked when odorants were presented are labeled in red and the response vectors during the after stimulus termination are shown in cyan. Results from three representative flies for each line/region is shown.

